# Illuminating the active virosphere with BONCAT and single virus genomic sequencing technologies

**DOI:** 10.1101/2025.08.05.666878

**Authors:** Maria Alvarez-Sanchez, Francisco Martinez-Hernandez, Aitana Llorenç Vicedo, Marina Vila-Nistal, Alon Philosof, Aditi K. Narayanan, Jamie C. Tijerina, Manuel Martinez-Garcia, Victoria J Orphan

## Abstract

Marine viruses impact biogeochemical cycles through cell lysis, releasing organic matter and nutrients that fuel ocean productivity. Identifying and quantifying the specific viruses active in these processes remains a priority in the field. Here, we introduce a click-chemistry method to fluorescently label, sort, and sequence the genomes of newly produced viral particles released from transcriptionally active host microbial cells, alongside the analysis of co-occurring inactive cells and viruses in environmental samples. This approach, called viral BONCAT-FACS, combines biorthogonal non-canonical amino acid tagging (BONCAT) with environmental sample incubation, followed by single-virus and single-cell sorting by flow cytometry (FACS). Genomic analysis of translationally-active cells and new viral progeny in coastal seawater incubations confirmed BONCAT labeling and successful sorting of diverse marine bacteria, microeukaryotic cells, and virioplankton, with stark differences in the predicted turnover of specific groups of infecting viruses, including Pelagiphages, Methylophages, a Flavobacteriales-associated novel “Far-T4” clade, algae-infecting giant NCLDV viruses, and parasitic virophages. Sequenced BONCAT-active cells showed a strong enrichment in viral contigs relative to the inactive cell fraction, suggestive of a large proportion of translationally-active virocells. This study illustrates the effectiveness of viral BONCAT-FACS for uncovering genome-resolved viral-host dynamics. By providing a direct approach for tracking active viral infections in natural environments, this method enhances our ability to investigate behavior and interactions of these nanoscale predators, expanding our understanding of their role in ecosystem dynamics.

## Introduction

Viruses and bacteriophages (herein referred to as viruses) are the most abundant biological entities in nature, exerting a significant impact across all ecosystems by influencing biogeochemical cycles^1, 2^, shaping genetic composition of host cells, and modulating microbial abundance through viral lysis^3–5^. After a successful viral infection, microbial cells (particulate organic matter, POM) are transformed into cell debris, releasing dissolved organic matter (DOM), that fuels ocean productivity. This process, known as the viral shunt^6^, recycles up to fifty percent of photosynthetically fixed carbon, an estimated 10 billion tons annually, in global ocean ecosystems^1, 4, 7^. During cell lysis, the viral progeny (the newly synthesized viral particles) are released into the environment, renewing the virosphere which is likely variable in both the rate of infection and burst size for different viral-host pairings (Fig. 1A). These factors, alongside the decay of viral particles, contribute to viral turnover^8^. Methodological limitations currently restrict our ability to quantify viral turnover and residence times at local scales, resulting in poorly resolved ecological dynamics^2, 9^.

**Figure 1.**
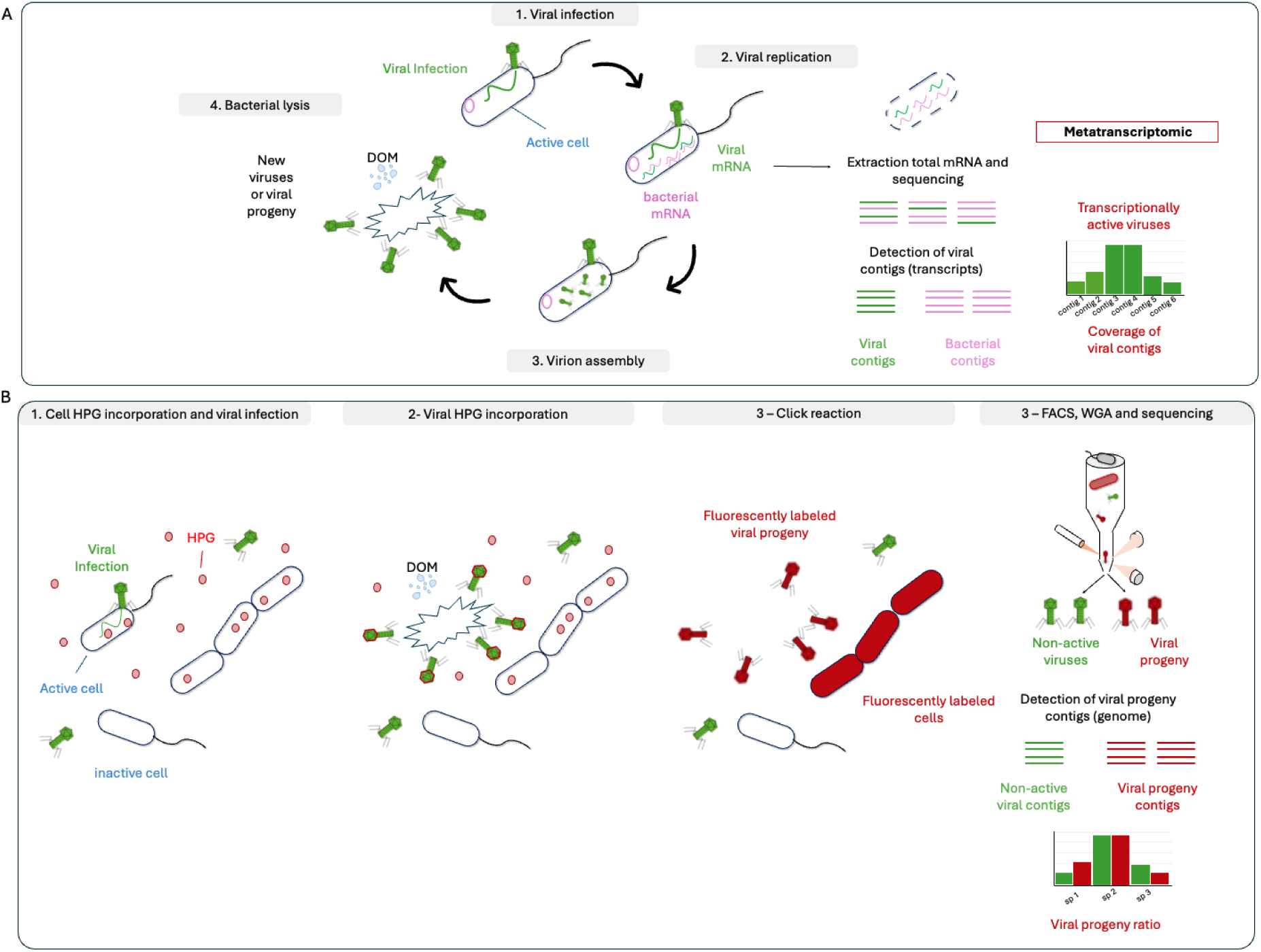
Schematic showing methodological steps of viral BONCAT-FACS and environmental metatranscriptomics for detecting active viral infections in uncultured microbial assemblages. A) Environmental metatranscriptomics allows the identification of viral transcripts within host cells that are produced during hijacking of the cellular replication machinery. The main steps during the lytic cycle represented. Here, the coverage of viral contigs is used to quantify the activity of different viruses present in the sample. However, after viral replication, only successful infections will culminate into the assembly of virions and their release into the ecosystem after cell lysis (viral progeny), contributing to cell mortality and the release of DOM. B) Schematic representation of the viral BONCAT process coupled with FACS and single-virus and cell-genomic technologies. Fluorescent labelling and sorting of the viral progeny with viral BONCAT-FACS allows identification of the genome of active virions produced in successful infections, allowing us to know the viral progeny ratio or turnover of virus populations and its contribution to the release of DOM in the ecosystem.

Traditional methods, such as environmental sample incubation with radiolabeled leucine or thymidine substrates^10^, has been used to assess viral production but these techniques lack information about the identity of the produced viral active progeny. Conversely, viromics and microbial metagenomics provide information about the total diversity, biogeography, relative abundance of viruses, and predicted viral-host interactions, but do not discriminate between the active newly produced virions and the resilient inactive free populations within the viral assemblage^11, 12^. Several studies have additionally employed metatranscriptomics to assess expression of genes indicative of viral infections within microbial cells as an estimate for viral lysis^13–19^ (Fig. 1A). This approach can be challenging however, as the proportion of viral reads in cellular metatranscriptomic libraries is low, and often this data cannot differentiate between successful lytic viral production versus abortive infections that do not result in cell mortality and new viral particles. Other methodologies employing DNA-based stable isotope probing from concentrated viral-like particles in sample incubations have also shown promise for tracking active viral infections^20–22^, however this technique requires large sample volumes and does not enable tracking at the level of individual active viral particles. The current limitations in methodologies for directly identifying and quantifying newly produced viruses among total viral assemblages in the environment represents a bottleneck to advancing our understanding of critical ecological processes such as cell mortality, viral-host dynamics, and viral-mediated nutrient cycling.

To address this shortcoming, we developed a synergistic methodological approach that combines protein targeted biorthogonal noncanonical amino acid tagging (BONCAT) for fluorescent labeling of newly produced viral progeny (viral-BONCAT)^23, 24^ with viral particle sorting by flow cytometry (FACS)^25, 26^ and single virus genome sequencing technologies^11, 27, 28^. Employing the viral BONCAT-FACS method in tandem with earlier developed cellular BONCAT-FACS, we demonstrate the detection, quantification, and genomic characterization of diverse viral progeny from successful active viral infections in environmental incubations, differentiated from co-occurring non-active (BONCAT-negative) viruses and microbial cells. The BONCAT assay requires sample incubation with an azide or alkyne-modified methionine analogs L-azidohomoalanine (AHA) or L-homopropargyl glycine (HPG) added at low concentrations. This bioorthogonal amino acid has been shown to be readily incorporated by diverse microorganisms (archaea, bacteria, and microeukaryotes)^29^ during protein synthesis using the cell’s native translational machinery, and AHA or HPG-labeled proteins are subsequently conjugated to a fluorophore using Cu(I)-catalyzed azide-alkyne cycloaddition (i.e. click-chemistry; Dietrich et al., 2006^23^). Applications of BONCAT (and BONCAT-FACS) in environmental microbial ecology studies (e.g. Hatzenpichler et al., 2014^29^), have provided insights into the physiology and genomic diversity of active uncultured members of microbial communities from diverse ecosystems, ranging from deep ocean sediments to hot springs, using incubation conditions mimicking the native environment, or used to track the activity of select community members in response to a stimulus^26, 29–37^. BONCAT has also been shown to label newly produced free virions after infection of a translationally active microbial host using epifluorescence microscopy^24^ as well as used to detect labeled viral proteins with proteomics methods in defined co-cultures^24, 38^. These studies highlight the potential of this protein-based click-chemistry labeling approach for studying virus ecology. Here we advance our earlier developed microscopy-based viral-BONCAT method for use with higher throughput flow cytometry combined with single virus sequencing technologies. With viral BONCAT-FACS, we demonstrate the ability to simultaneously sort and genomically characterize discrete populations of newly produced viruses and co-occurring active and inactive microbial cells spanning large size range and fluorescence intensities in the sample. This methodological approach assists with deconvolving complex viral-host interactions in environmental samples, discriminating between active (lytic) viral clades with likely ecological impact, as opposed to inactive viral populations with slow decay rates. Importantly, as viral BONCAT-FACS is based on labeled proteins (e.g. capsids) rather than nucleic acids, this methodological approach should in principle be able to capture both DNA and RNA viruses and can identify them independent of whether related genome sequences are previously known. We demonstrate the efficacy of the viral BONCAT-FACS method coupled with single-virus and cell sequencing technologies for investigating complex communities in environmental samples, here characterizing paired active and inactive dsDNA viruses and microbial communities in coastal seawater incubations from the Pacific Ocean and Mediterranean Sea.

## Results

### Optimizing viral-BONCAT for flow cytometry sorting of environmental viruses

In the viral-BONCAT assay, newly produced viruses are labeled during infection of an anabolically active (i.e. BONCAT-positive) microbial host, where the methionine analog AHA or HPG is directly incorporated into viral proteins derived from the host during the lytic cycle. Subsequently, free nanometer-sized viral progeny released after cell lysis can be directly visualized and identified by fluorescence labeling (Fig. 1B, Supplementary Fig. 1).

The fluorescence intensity of BONCAT-positive environmental viruses using the original viral-BONCAT protocol described in Pasulka et al^24^ was inadequate for FACS detection and sorting, as small particle detection by flow cytometry requires a stronger fluorescence signal compared with epifluorescence microscopy. Optimizing the click-chemistry protocol, we boosted the fluorescent labeling and demonstrated the ability to consistently detect of viral-like particles (VLPs) above the background noise during flow cytometry sorting at two independent sorting facilities, illustrating its transferability between labs (Fig. 2, Supplementary Fig. 2)^11, 27, 28^. Specific protocol modifications for BONCAT included performing the click reaction under oxygen-free conditions using argon gas, as the copper (I) catalyzed click reaction is sensitive to oxygen^39^ and dampens the fluorescence signal (detailed in methods; Supplementary Fig. 1). Additionally, the click reaction time was extended from the standard 30-minute incubation to 6 hours, which also substantially enhanced the fluorescence signal from individual viral-like particles (Fig. 2).

**Figure 2.**
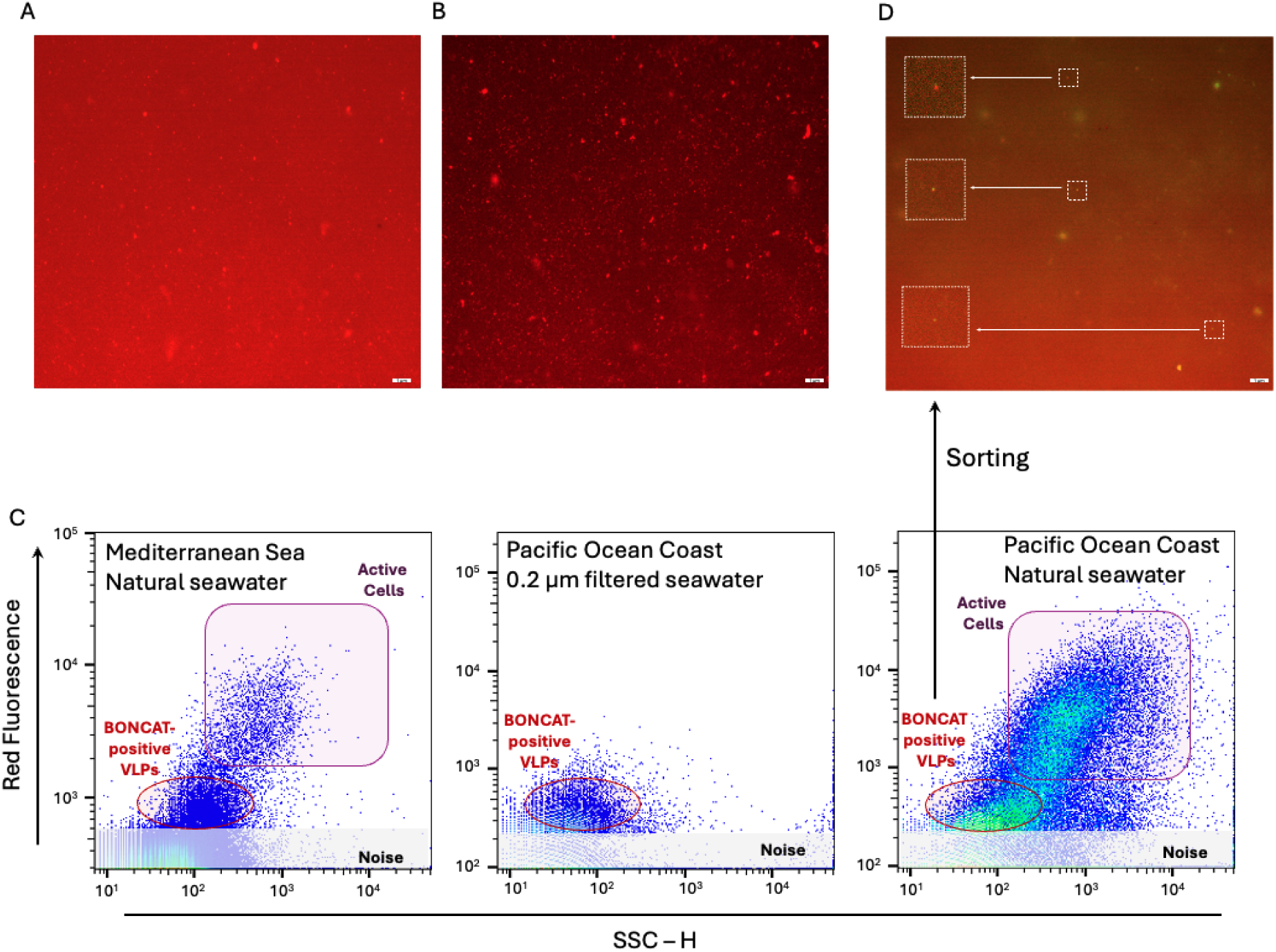
Viral BONCAT-FACS applied to study active viruses in coastal seawater samples from the Mediterranean Sea and California Coast. A) Fluorescence microscopy image of active viruses (viral progeny) from a coastal Mediterranean seawater sample after a 30 min CLICK reaction in the viral BONCAT protocol using AF647-Pycolyl-Azide (red fluorescence) B) Effect of increasing the click reaction to 6 hours using the same viral BONCAT protocol. C) Flow cytometry plots showing the positive active cell and viral populations from coastal seawater samples collected in the Mediterranean Sea and California coast. SSC-H in x-axis side scatter (relative units); Red fluorescence in Y-axis from AF647-Pycolyl-Azide (relative units). D) Imaging of BONCAT-labeled VLP’s by epifluorescence microscopy from the sorted active viral fraction. Magnifications of representative viruses that were stained with both AF647-Pycolyl-Azide (red) and SYBR-Gold (green) are highlighted in the boxes. The scale bar in panels B, C, and E is 1 μm.

Freshly collected coastal surface seawater samples (0.25 to 2 L) from the Mediterranean and the Pacific were amended with 10 μM HPG and incubated for 5 days under temperature and diel light conditions similar to that in the environment (detailed in methods) allowing for the assimilation of HPG into proteins within translationally active microbial cells and their infecting viruses^29^ (Fig. 1, Supplementary Fig 1).

From these two independent coastal seawater experiments, separate pools of positive BONCAT-labeled virions and cells, along with the co-occurring non-active cell and viral fractions (BONCAT negative) were sorted, whole-genome amplified^11, 28^, and Illumina sequenced. From the Mediterranean Sea sample, 41,000 viral-like particles (VLPs) from the viral progeny assemblage (BONCAT positive VLPs) and 3,100 active microbial cells were sorted and sequenced. From the coastal Pacific Ocean BONCAT incubation, 100,000 VLPs from the viral progeny and 50,000 non-active VLPs were sorted and sequenced in addition to 25,000 BONCAT active and non-active microbial cells. Detection and corroboration of *bona fide* viral contigs from these sequencing datasets was performed using a combination of Virsorter 2.0^40^, geNomad^41^, and compared against the IMG/VR^42^ database v4.1 (using a protein similarity threshold of 80% and an alignment coverage of 90%; Fig. 3). Sequence datasets of the BONCAT-labeled viral fractions showed minimal contamination from cellular debris or outer membrane lipid vesicles, which have been shown to be common in seawater and are difficult to distinguish from viruses based on size and nucleic acid staining alone^43–46^. Epifluorescence microscopy was also used to independently confirm that the sorted viral fraction indeed contained BONCAT-tagged and SYBR stained VLPs (Fig. 2D).

**Figure 3.**
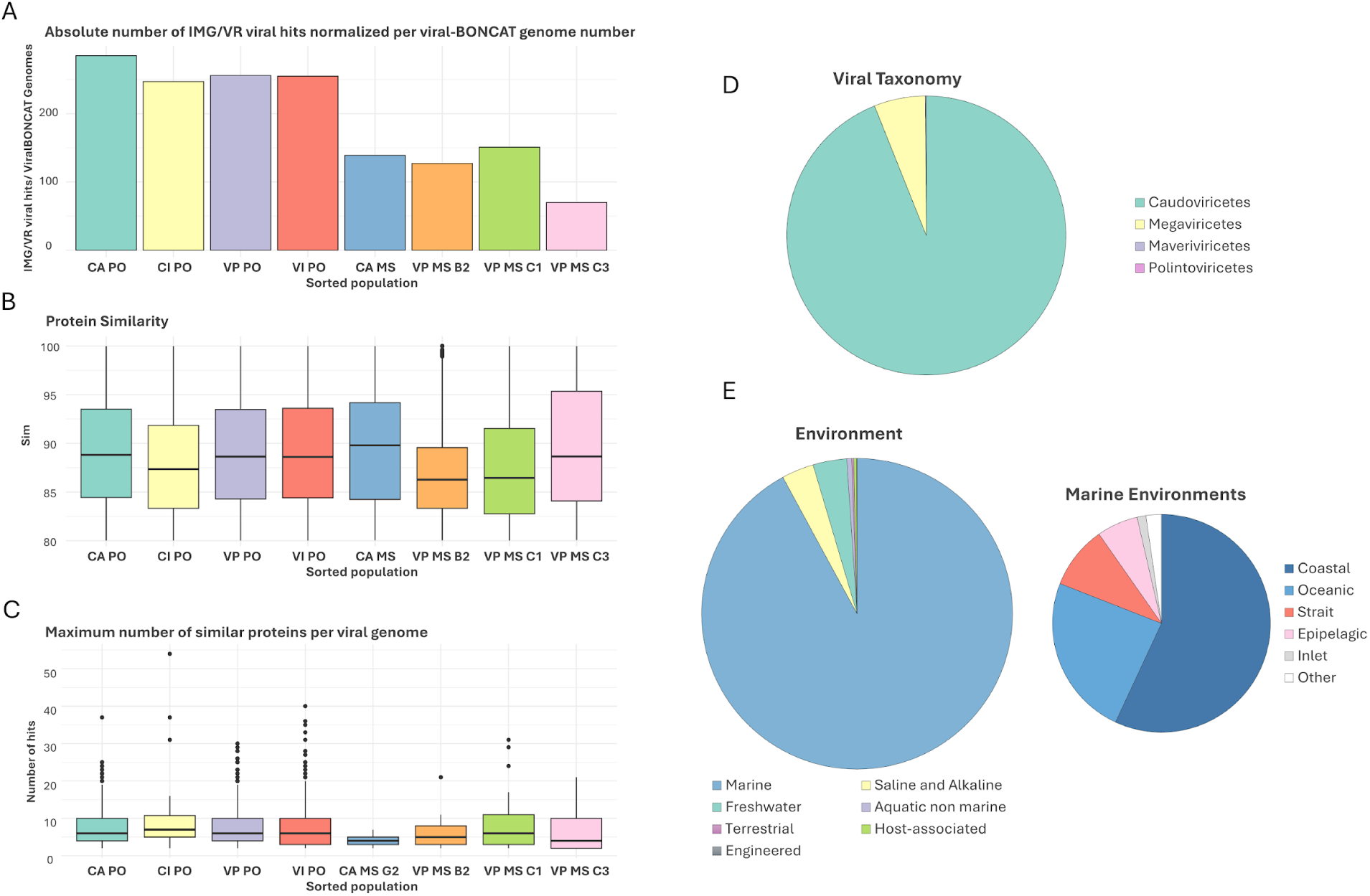
Sequencing results and genomic analysis of sorted viruses obtained by viral BONCAT-FACS. A protein blast against curated viral genomes identified with high confidence was performed to confirm the viral origin of the recovered viral BONCAT sequences after flow cytometry sorting of BONCAT positive cells and viruses and corresponding inactive fractions from the Pacific Ocean (PO) and Mediterranean Sea (MS). Panel A shows the mean value of protein hits for each contig from the viral BONCAT sorts for the following sorted fractions-CA: Cells Active, CI: Cells Inactive, VP: Viral Progeny, VI: Viruses Inactive. Panel B shows the mean amino acid similarity of these hits, and panel C represents the maximum number of similar proteins for the viral BONCAT contigs against reference viral genomes. Panel D shows the proportion of recovered viruses belonging to major taxonomic groups assigned by geNomad. Panel E summarizes the source environment for the closest matches to reference viral genomes.

### Genomic characterization of BONCAT active and inactive virioplankton and lineage specific variation in viral production

Across all sorted cellular and viral samples including both active and non-active BONCAT faction, we identified a total 2,192 viral contigs greater than 1.5 Kb (Fig. 3, Table 1). Of these, 74% of contigs (n=1,630) contained two or more viral hits in the comparative analysis against the viral IMG/VR v4.1 database. Remarkably, nearly half of these viral contigs (n=803) were detected in the BONCAT-positive viral fraction. This confirmed the successful recovery of BONCAT-labeled environmental viruses by FACS and demonstrated the feasibility of our overall workflow of viral-BONCAT combined with FACS and single-virus sequencing^11, 27, 28, 47^. The population structure and diversity of BONCAT-positive and non-active (BONCAT-negative) viral contigs >5 Kb length (n=1,223) was analyzed by generating a protein sharing network using vConTACT2^48^ v.0.9.19 (Fig. 4). Similar viral genomes were recovered from both the sorted viruses infecting translationally active hosts and the non-active viral assemblage. However, different viral clusters within the network displayed large variations in the relative proportion of active and non-active viruses, which we defined here as the viral progeny ratio (i.e., normalized proportion of newly synthesized viral progeny vs. non-active viral contigs; see Methods, Supplementary Table 1 and Supplementary Data 1; Fig. 4). This comparative analysis identified population-specific differences in viral activity and turnover rates among commonly recovered marine viruses. For example, viruses related to the uncultured pelagiphage vSAG 37-F6^11, 49^ (average amino acid similarity of ≈75%, Supplementary Data 2), one of the most abundant phages in the oceans^11, 49^, were associated with a viral cluster that was dominated by newly produced, BONCAT-positive viruses (viral progeny ratio = 78.95%, n=17; Fig. 4, Supplementary Table 1). Similarly, a second dominant viral cluster also stood out for being almost exclusively comprised of active viruses (viral progeny ratio= 82.57%, n=199; Fig. 4, Supplementary Table 1). This cluster contained several newly described uncultured Far T4-like phages from marine environments^50^, and an isolated *Rhodothermus* phage RM378^51^ as the only reference genome. Phylogenetic analysis of the major capsid protein, Gp23, and other core proteins (large terminase, prohead, and portal proteins) from several of our recovered viral contigs in this cluster confirmed relatedness to these uncultured Far-T4 phages (Fig. 5; n=15). The majority of the recovered Far-T4-like phage genomes (12 out of 15) came from the BONCAT-positive viral fraction, suggesting fast turnover of these viruses over the incubation period. Using the tool iPHoP^52^, we screened for potential microbial hosts of the Far-T4-like phages. Several diverse host lineages were identified, with the largest fraction (113 out of 235, or 48%) belonging to the order Flavobacteriales (see Supplementary Data 3). Consistent with this finding, Flavobacteriales genome sequences were 15 times more abundant in the BONCAT-active cell fraction (3% of the total bacterial reads) relative to the non-active sorted cells (0.2%) in the corresponding FACS-sorted microbial cell fractions (Supplementary Fig. 3; data available in Supplementary Data 4). The proportion of putative microbial hosts that were BONCAT-positive (active cells) in the seawater incubation, in tandem with the predominance of Far-T4-like phages in the viral progeny, is suggestive of *kill-the-winner* dynamics, resulting in fast turnover of this active group of viruses^53, 54^. While this current study focused on a single timepoint, future work using viral BONCAT-FACS for time course experiments would enable longitudinal tracking of active and inactive cell and virus populations, contributing to our understanding of host-virus dynamics.

**Figure 4.**
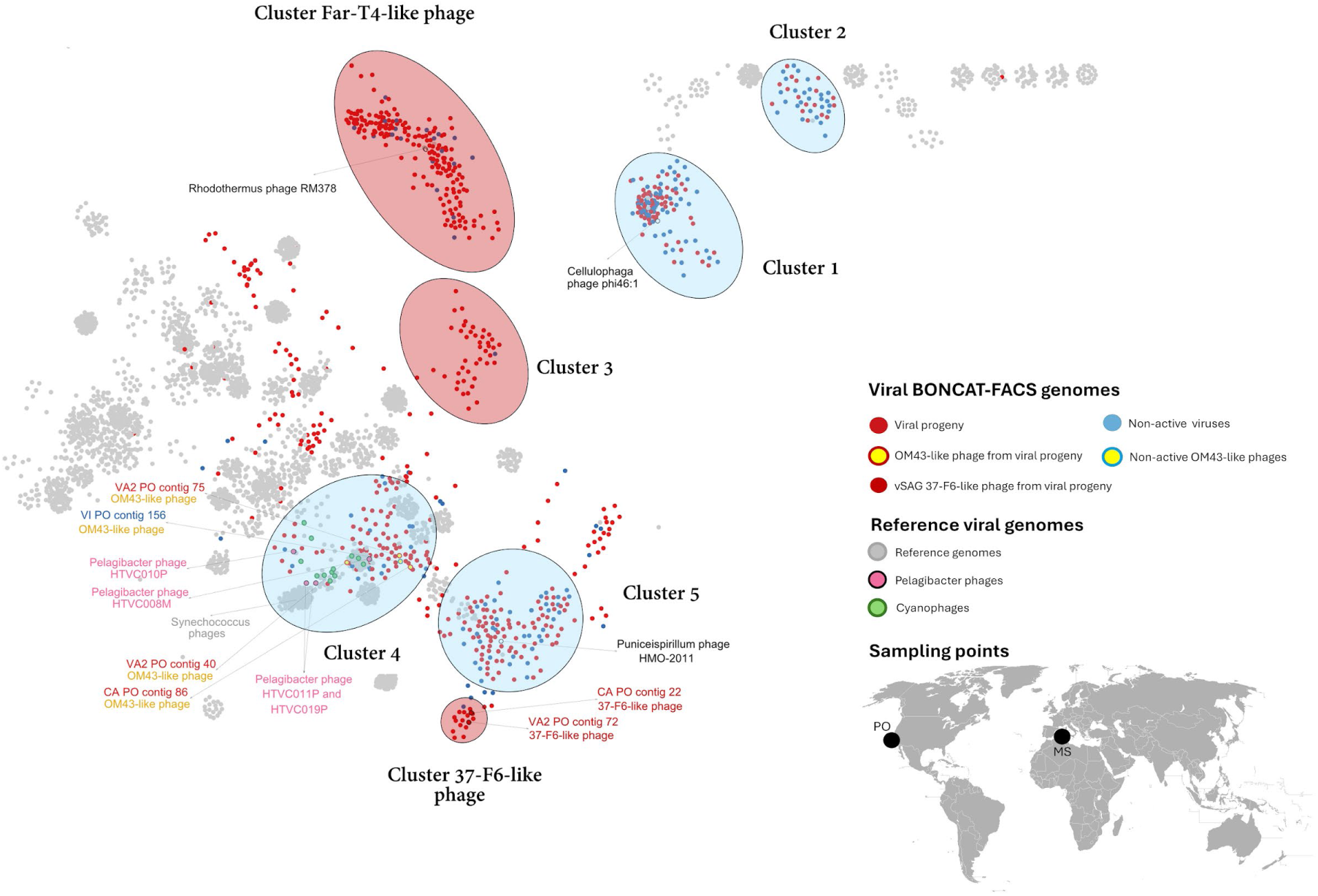
Protein cluster network of viral genome fragments obtained by viral BONCAT-FACS. Viral genomes >5 Kb obtained from coastal Mediterranean Sea (MS) and Pacific Ocean (PO) seawater incubations (inset) were clustered based on their protein similarity. Viral populations among distinct clusters showed different levels of activity and putative turnover rate based on the number of BONCAT active (red circles) and non-active (blue circles) viruses recovered from the incubations. Grey points represent reference viral genomes from the *Prokaryotic Viral RefSeq 201* database. Red ellipses depict clusters dominated by viruses from the viral progeny fraction (BONCAT positive) with little to no representation of these groups in the inactive fraction, reflective of high activity and fast turnover rate. Conversely, blue ellipses highlight viral clusters with minimal viruses from viral progeny compared to non-active, likely representing viral populations with lower lytic activity and slower turnover rates. Select reference genomes from isolated phages are highlighted and labeled.

**Figure 5.**
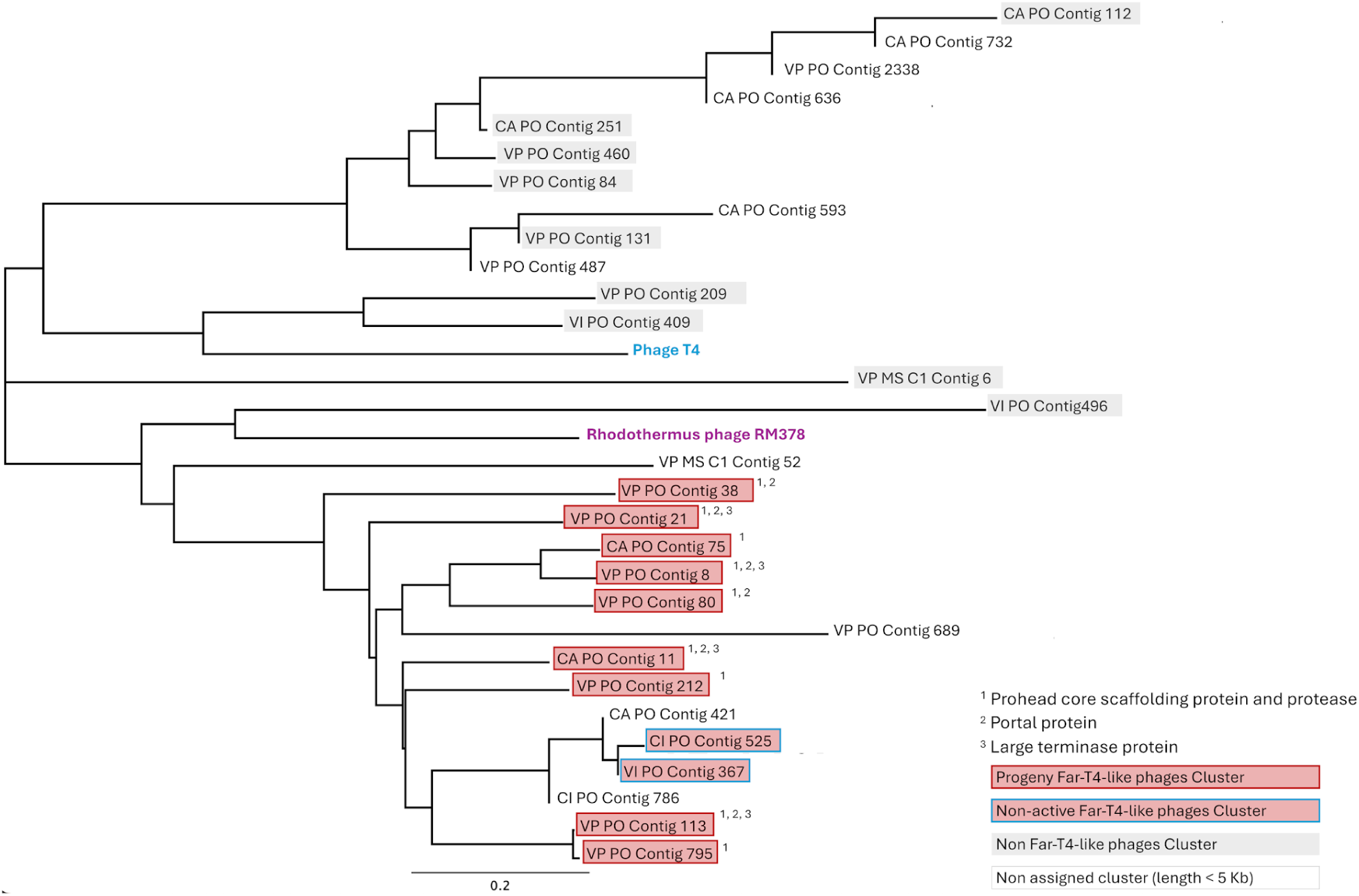
Phylogenetic tree of the structural viral protein gp23 used as marker for the T4-like phages. Red labels indicate viruses belonging to the Far-T4-like cluster depicted in Fig. 3. Grey labels represent viral contigs contained in other clusters with a gp23-like structural protein. Contigs with no labels are viral BONCAT contigs smaller than 5 Kb that were not considered in the protein viral network. The superscripts indicate those viral genomes recovered in our study sharing core proteins with Far-T4 phages.

**Table 1.**
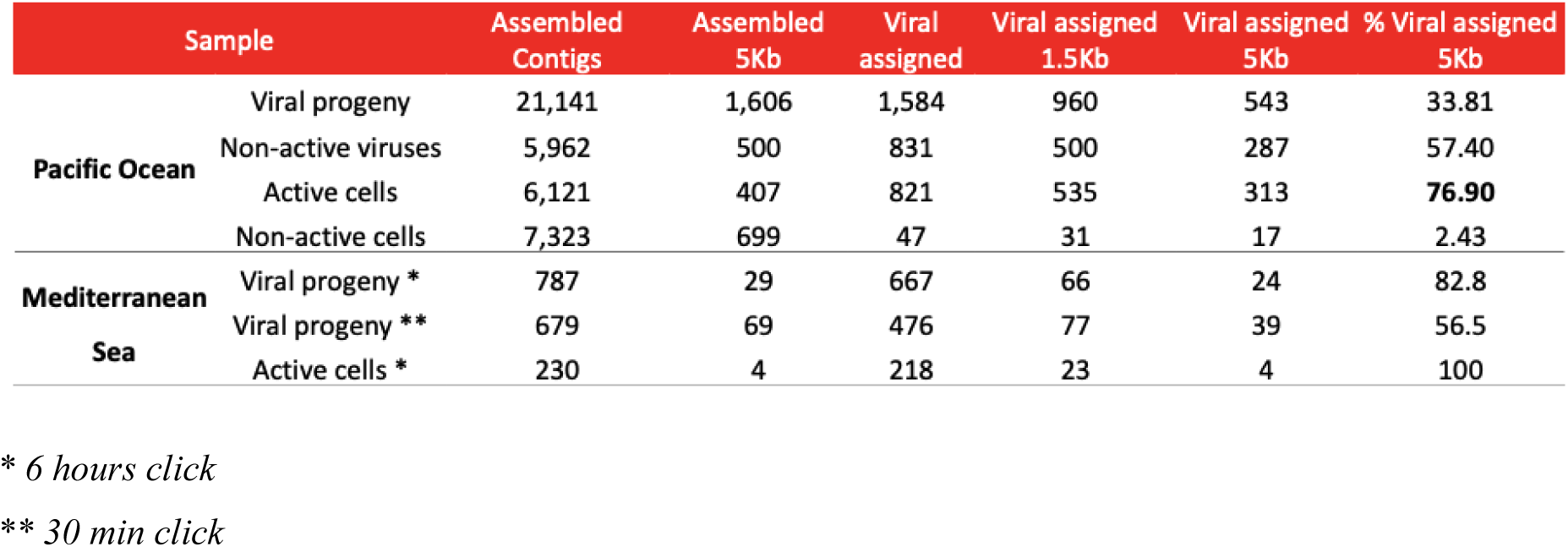
Assembled viral contigs assigned as viruses in each of the sorted populations.

### Analysis of viral contigs in the BONCAT-FACS sorted active and inactive microbial cells and linking active viruses and hosts

In addition to the information gained from direct sequencing of the sorted viral fraction, the analysis of the co-occurring microbial cells from the BONCAT-active and inactive fractions detected a greater proportion of viral contigs in the translationally active cells than inactive ones in our seawater incubations. These viral contigs comprised approximately 77% of the total assembled contigs greater than 5 Kb in the BONCAT-active microbial cell fraction, largely associated with the Caudoviricetes (Table 1; Fig. 3D). In contrast, the sorted inactive microbial fraction from the same sample (i.e., BONCAT-negative cells) contained only 2.4% viral contigs. This data suggests a higher infection rate and viral pressure among the translationally active members of the microbial community.

The taxonomic composition of the sorted BONCAT-active and inactive microbial cell fraction was also notably different. While the active microbial cell fraction sequenced from both field sites showed an abundance of *Pseudomonodota* (formerly Proteobacteria) representing ∼72-75% of the bacterial reads, the inactive cell fraction (coastal Pacific site) instead was largely associated with members of the Planctomycetaceae, representing ∼28% of bacterial reads, with less than 0.1% in the active fraction (Supplementary Fig. 3; Supplementary Data 5-6). Among the BONCAT-sorted *Pseudomonodota*, members of the alpha- and betaproteobacteria were dominant and represented largely by the common marine lineages Pelagibacteriales (SAR 11) and Nitrosomonadales. Within the Nitrosomonadales, several taxa were affiliated with the methylotrophic clade OM43^55^, representing 2% and 17% of the assigned *Pseudomonodota* reads from Pacific Ocean and Mediterranean Sea, respectively (Supplementary Fig. 3; Supplementary Data 4 and 7). Described members of the OM43 clade are submicron-sized cells with streamlined genomes that are known for their ability to utilize C1 compounds and are common in coastal environments^55, 56^.

A deeper characterization of OM43 diversity among the sequenced active and non-active cell fractions revealed the prevalence of a specific strain, *Methylophilales* sp. H5P1^57^, from the coastal Pacific Ocean site (Supplementary Fig. 4). Corresponding with recovery of this bacterial strain, four of the assembled viral contigs from our paired sorted viral fraction were closely related to the previously characterized methylophage group Melnitz, which has been previously shown to infect strain H5P1^58^ (Fig. 4; Supplementary Data 8). These data are further evidence for the effectiveness of coupled cell- and viral-BONCAT as a tool for assessing viral-host pairings. Melnitz myophages have been reported from other temperate coastal and subtropical gyre environments^58^. Our four assembled methylophages were co-located in a cluster with pelagiphages and cyanophages in the protein viral network (Cluster 4, Fig. 4; more detailed in Supplementary Data 1), a feature also reported in the previously isolated Melnitz phages^58^. Notably, these groups represent viral clades previously reported to have low production rates in the ocean^13, 59^ and correspondingly, our BONCAT-estimated relative progeny ratio for the Melnitz-like phages group (Cluster 4) was 65.96% (n=78; Fig. 4 and Supplementary Table 1), indicating lower turnover rates compared with other co-occurring phages like the Far-T4-like viral cluster.

### Viral BONCAT-FACS recovery of eukaryotic viruses and parasitic virophages

Alongside the recovery of diverse bacteriophages, the viral BONCAT active fraction also recovered several eukaryotic viruses affiliated with nucleocytoplasmic large DNA viruses (Nucleocytoviricota or NCLDVs)^60^. A total of 12 viral contigs (>5 kb) associated with Phycodnaviruses, likely infecting algae, were recovered from the active virioplankton fraction from the coastal Pacific, with an additional four Megaviricetes viral contigs recovered in the Mediterranean Sea dataset. Within the Pacific Ocean samples, nine additional Phycodnavirus contigs were detected from the non-active viral fractions. Phycodnaviruses (n=11) were also recovered from the active microbial cell fraction, indicating active infection of microeukaryotes (presumably a marine alga) over the course of the incubation (Supplementary Fig. 5, Supplementary Table 2). Interestingly, alongside the Phycodnaviruses, genomes from two virophages (Lavidaviridae family) were also recovered in the same sorted BONCAT-positive viral fraction from the coastal Pacific (Supplementary Table 3). Virophages are parasitic phages that infect NCLDV viruses^61^ and detection of an ongoing ‘superinfection’ with viral BONCAT-FACS opens up new opportunities for studying infection dynamics in environmental samples for a broad diversity of bacteriophages and eukaryotic viruses, alike.

## Discussion

In this study, we combined high-throughput BONCAT-FACS with advanced low input viral genome sequencing technologies to investigate the active fraction of virioplankton, the smallest and most diverse biological entities in the ocean. Viral BONCAT-FACS provides a targeted method to differentiate and sort newly produced viruses and their microbial hosts based on translational activity, enabling genomic characterization of both BONCAT-labeled and unlabeled components from the same sample. This represents a unique and complementary approach to community-wide methods in viral ecology such as metagenomics^62^, transcriptomics^13–19^, stable isotope probing^20, 63^, and radiotracer techniques^10^, which lack the resolution to identify individual active virions or infected cells. The integration of activity-based, single-cell and virus sorting with genomic analysis of the active and inactive fractions, yields information on often hidden dynamics within populations of viruses and their hosts, representing an important step towards developing a systems-level understanding of microbial interactions and activity within ecosystems. These datasets, combined with time series studies, can help refine ecological models of viral production, turnover, and their functions in regional and global biogeochemical cycles^64, 65^.

The viral-BONCAT method was initially developed as a fluorescence microscopy assay to visualize and quantify newly produced viruses in model microbial systems and seawater incubations^24, 38^. We adapted and optimized the BONCAT protocol for high throughput flow cytometry sorting and single virus sequencing, enhancing fluorescence signal intensity to reliably detect, sort, and genomically characterize nanometer-scale viral particles and host microbial cells concurrently. The transition to flow cytometry, however, required overcoming technical limitations, as these submicron entities are at the theoretical limit of detection for most flow cytometers^11^. With this optimized method, we sorted both BONCAT-active and inactive microbial cells alongside viral particles from coastal seawater samples from the Mediterranean and Southern California and demonstrated its utility for resolving complex virus host dynamics in the environment. Our recovery of OM43 infecting Melnitz-like phages from the same incubation as their previously described methylotrophic host, Methylophilales sp. H5P1^58^ for example, provided confirmation of the ability of the viral BONCAT-FACS method to detect active infections and link viruses with their putative hosts within diverse environmental communities.

Across both datasets, our results revealed notable variation in viral production across different lineages. High turnover was observed for specific SAR11-associated pelagiphages (e.g. vSAG 37-F6-like phages) and the recently described Far-T4-like phages which we putatively linked to Flavobacteriales hosts, with high enrichment for both in the BONCAT-positive viral fraction. The recovery of BONCAT-positive phages belonging to the vSAG 37-F6 group was particularly noteworthy as this uncultured pelagiphage group, originally characterized by single-virus genomics methods, was reported to be one of the most abundant and ubiquitous viruses in the marine virosphere^11, 27, 49, 66^. This viral BONCAT-FACS generated data is consistent with previous reports from whole community viral metatranscriptomes from coastal environments which reported the high expression of vSAG 37-F6 transcripts relative to other pelagiphages, such as HTVC010P or HTVC011P, indicating a higher rate of host infection^13^. Our results expand upon these earlier findings, providing independent support for higher infection rates and turnover by the lytic vSAG 37-F6 group in coastal seawater among the co-occurring pelagiphages in our incubations. Given its ubiquity and activity in ocean ecosystems, this specific phage lineage is likely to be an important contributor to nutrient recycling through the viral shunt^67^ and warrants further study (Fig. 4).

Our BONCAT-based estimate of lower viral production and turnover for other lineages related to cyanophages, methylophages, and the aforementioned HTVC010P and HTVC008M pelagiphages (e.g. Cluster 4, Fig. 4; Supplementary Table 1) was also consistent with their potential targeting of slower growing bacterial hosts, as has been observed in earlier studies^13, 59^. Other viral groups in the protein cluster network had even lower turnover rates, such as the phages in Cluster 1 likely infecting members of the flavobacteriales, including *Cellulophaga* (Fig. 4, Supplementary Table 1 and Supplementary Data 1). This slow predicted turnover was distinct from the high progeny ratio calculated for the flavobacteriales-associated Far T4-like cluster and, as observed among different pelagiphage lineages, highlights the variation in viral pressure among related host bacteria in coastal zones. Collectively, these lineage-specific patterns support ecological models such as *kill the winner*^53, 68^, where rapidly growing microbial host populations are impacted by elevated viral pressure and rapid phage production, while other phage lineages persist with comparatively lower production rates and turnover.

Beyond the dynamics detected among different phage lineages by viral BONCAT-FACS, the concurrent sorting and analysis of BONCAT-active and inactive microorganisms unexpectedly revealed substantial enrichment in viral contigs associated with the active cellular fraction (77%) relative to the inactive cells (2.5%) in the same incubation, suggesting that a high proportion of the translationally active planktonic bacterial community may be virocells^69–72^. The relationship between (presumed) viral infection and microbial activity is consistent with earlier reports demonstrating a positive correlation between total viral abundance and bulk bacterioplankton productivity^73^ however the percentage of virocells in our BONCAT-FACS dataset is substantially higher than reported values other FACS-enabled studies of bacterioplankton communities (∼34% of cells)^12, 74^. This large discrepancy may be attributed to the fact that in these earlier studies, total cells were sorted rather than differentiated based on translational activity.

As a positive BONCAT signal requires cellular uptake and incorporation of the methionine surrogate during protein synthesis, our findings suggest that microbial cells continue to actively take up nutrients just prior to or during viral infection. This interpretation aligns with prior stable isotope probing experiments in which *Synechococcus* infected by cyanophage continued to assimilate exogeneous nitrogen and synthesize proteins throughout the infection cycle^75^. Should this pattern hold across systems, the ecophysiological interpretation of BONCAT activity may need to be broadened to reflect not just cellular anabolic activity, but also the presence of viral infection.

The microbe-virus interactions recovered in our viral BONCAT-FACS experiments were not limited to bacteria and their phages but also included several giant viruses known to infect microeukaryotes (NCLDV viruses belonging to the Nucleocytoviricota), highlighting the technique’s ability to capture a broad diversity of active microbes and lytic viruses. The simultaneous activity-based sorting of viruses and cells by FACS offers an important methodological advantage by directly capturing newly produced viral progeny across a broad size range, avoiding the size-selective bias that often excludes giant viruses from filtered submicron viral concentrates. We additionally detected a potential virophage from the *Lavidaviridae* group within the active viral fraction, indicating there was active production of these virophages during the incubation (Supplementary Table 3). *Lavidaviridae* are known to parasitize giant Nucleocytoviricota viruses, however the specific association in this case remains unresolved due to the lack of a confirmed eukaryotic host. NCLDV and virophages are widely distributed in ocean ecosystems^61, 76–79, 80^ and the nature of this superparasitism and its impact on host ecology remain areas of active study. Future investigations that integrate the unique perspective of single-cell and virus-targeted BONCAT-FACS with established viral ecology methods have the potential to expand our understanding of their ecology and the underlying interactions between microeukaryotes, viruses, and their parasites in nature.

While viral BONCAT-FACS yields unique insights, several methodological considerations warrant discussion. As with all incubation-based methods, BONCAT experiments may be subject to potential artifacts such as bottle effects which can alter community composition and activity. Additionally, variability in HPG sensitivity and assimilation among microbial taxa may influence BONCAT results^81^. These factors should be independently evaluated when applying viral-BONCAT to new environments. In the context of our study, the use of BONCAT in marine planktonic communities has been previously assessed and found to yield results comparable with other methods. Prior investigations with coastal seawater for example showed consistent results between BONCAT-based assays and independent measures of community productivity and viral production^24, 82^. Additionally, the low concentration of HPG used in our incubation experiments was below that reported to impact the growth of some cyanobacteria^81^. Despite the potential for perturbation, the application of viral BONCAT-FACS was able to successfully recover and genomically characterize newly produced marine viruses and diverse translationally active microbial hosts from coastal seawater. With appropriate optimization, this approach can be applied to study viral-host dynamics in diverse ecosystems. This activity-based sorting of microbes and viral progeny complements existing meta’omics methods and introduces a valuable approach for detecting active infection and lineage specific estimates of viral turnover within the context of complex microbial communities in nature.

### Conclusions

The development and successful application of viral BONCAT-FACS represents an important advancement in viral ecology complementing existing ‘omics-based methods. This activity-based sorting and genomic-sequencing approach enables the differentiation of newly produced viruses and their actively infected microbial hosts from co-occurring inactive populations in natural communities. Our initial application of viral BONCAT-FACS yielded insights into diverse marine virus-host interactions in coastal ecosystems, with lineage-specific differences in turnover predicted for viruses infecting common members of the bacterioplankton including Flavobacteriales, SAR11, OM43, Cellulophaga, and eukaryotic phototrophs. The unexpected finding of a high prevalence of virocells among translationally active members of microbial communities necessitates a reconsideration of traditional interpretations of microbial activity and highlights new directions for exploring the impacts of viral infection and its influence on nutrient cycles. Looking forward, viral BONCAT-FACS holds promise for revealing previously hidden viral-host interactions across diverse ecosystems, from deep-ocean sediments to industrial bioreactors and animal and plant-associated microbiomes, expanding our systems-level understanding of microbial communities and their ecological and biogeochemical significance.

## Data availability

The raw reads from the active and non-active cell and viruses are available in the Zenodo repository: https://doi.org/10.5281/zenodo.12903744 (Pacific Ocean Samples) https://doi.org/10.5281/zenodo.13149427 (Mediterranean Sea samples)

## Acknowledgements

This work was funded by the U.S. Department of Energy, Office of Science, Office of Biological and Environmental Research under Award Number DE-SC0022991 and subaward No. *S591062,* with additional support from the Spanish Ministry of Universities (ref. *MARSALAS21-15*), Agencia Estatal de Investigación (PID2021-125175OB-I00) Generalitat Valenciana (ref. *APOSTD/2020/237*), the Simons foundation PriME collaboration, and an investigator grant from the NOMIS foundation (to VJO).

## Disclaimer

This report was prepared as an account of work sponsored by an agency of the United States Government. Neither the United States Government nor any agency thereof, nor any of their employees, makes any warranty, express or implied, or assumes any legal liability or responsibility for the accuracy, completeness, or usefulness of any information, apparatus, product, or process disclosed, or represents that its use would not infringe privately owned rights. Reference herein to any specific commercial product, process, or service by trade name, trademark, manufacturer, or otherwise does not necessarily constitute or imply its endorsement, recommendation, or favoring by the United States Government or any agency thereof. The views and opinions of authors expressed herein do not necessarily state or reflect those of the United States Government or any agency thereof.

### Author information

These authors contributed equally: Maria Alvarez-Sanchez, Francisco Martinez-Hernandez

## Contributions

V.J.O. and M.M.G., conceived, designed, coordinated the project, and provided funding

M.A.S., F.M.H., A.L.V., M.V.N., A.P., A.K.N., M.M.G., and J.C.T., performed experiments.

M.A.S., and F.M.H. analyzed the data.

M.A.S., F.M.H., M.M.G., and V.J.O. wrote the manuscript.

## Ethics declarations

Competing interest

The authors declare no competing financial interests.

