## Supplementary material for "Illuminating the active virosphere with BONCAT and single virus genomic sequencing technologies"

The authors declare no competing interests.

**This supplementary file contains a Method section, references, 6 figures and 3 tables. A separate Excel file titled “Supplementary Data 1–8.xlsx” includes eight supplementary datasets described in the section “Supplementary Data Legends” below.**

**METHODS**

**Sample collection and processing**

Two independent experiments were conducted using seawater samples from the California Pacific Coast and Mediterranean Sea. Pacific Coast seawater sample (0.50 L) was obtained from the Corona del Mar coast at Kerckhoff Marine Laboratory (California, USA, 33° 35′ 47.6″ N, 117° 52′ 48.9″ W) on April 29^th^, 2023. Mediterranean surface seawater sample (4.00 L) was collected from Cape the Huertas (Alicante coast, Spain, 38° 21′ 14.3″ N, 0° 25′ 36.6″ W) on November 9^th^, 2023. In both cases, seawater was transported to the laboratory as quickly as possible, within a time frame of less than 2 hours. A total volume of 0.25 L (Pacific coast) and 2.00 L (Mediterranean Sea) were incubated with 10 µM of L-Homopropargylglycine (HPG; amino acid analog of methionine) in Erlenmeyer flasks during 5 days at 18ºC and under light:dark photoperiods of 12h:12h (Mediterranean coast) and 14h:10h (Pacific coast), mimicking natural conditions. In parallel, for each sampling point, same volumes (0.25 and 2.00 L) of seawater were incubated without HPG under same conditions, used as control to discriminate the active and non-active populations during FACS after the click chemistry reaction. From the Mediterranean Sea sample, 0.25 L were employed to run a 30 min click reaction, while the rest of the volume (1.75 L) were used for a 6 h click reaction. From the Pacific coast seawater sample, all seawater volume was used for a 6 h click reaction. After the 5-days incubation, the seawater (with and without HPG) was concentrated up to an approximate volume of 15 mL using the tangential flow filtration system Vivaflow 30,000 MWCO PES cassettes (Vivaflow 200, Sartorius), and subsequently Amicon Ultra Centrifugal filters, 100 kDa (Amicon Ultra, Merck Millipore) were used to finally ultra-concentrate the sample by centrifugation (2,500 xg, 3 min) up to a final volume of 208 µl. Then, samples were washed with 5 ml of 0.02-μm-filtered 1× PBS, pH 7.4 and concentrated by centrifugation (2,500 x *g*, 3 min). This washing process was repeated 3 times. To reduce the viral decay and cell activity, samples were kept at 4ºC during all the concentration and washing processes.

**BONCAT-Click chemistry reaction stain of active populations**

Standard click reaction for fluorescently labeling incorporated HPG in active cells^1,2^ and viral progeny^3^ was optimized and performed in solution, without fixation, to allow the discrimination, sorting, and genome recovery of viral progeny and active cells using a combination of FACS and single-cell and -virus technologies. To minimize the oxygen concentration, click reaction was conducted in a PCR tube (250 µl) as follows: 208 µl of sample volume and 42 µl of click mix (see below). As blanks in BONCAT, 208 µl of 1X sterile PBS were used. The click mix is made in 2 steps by the dye-premix (0.5 mM CuSO_4_, 2.5 mM THPTA and the 0.1 µM dye, final concentrations) and the reaction buffer (25 mM Aminoguanidine, 25 mM Sodium Ascorbate final concentrations, and 1X PBS). The work solutions of 20 mM CuSO_4_ (dissolved in 0.02 µm filtered MilliQ water), 100 mM Aminoguanidine and 100 mM Sodium Ascorbate (both dissolved in 0.02 µm filtered 1X PBS) were prepared freshly just before starting the click reaction. The 50 mM THPTA, the 10 µM azide-modified dye (AF647-Picolyl-Azide) stock solutions were previously diluted in 0.02 µm filtered MilliQ water and DMSO, respectively, and they were frozen at -20 ºC until use. To remove the dissolved oxygen, immediately before the preparation of the click mix, all the reagents and the 0.02 µm filtered 1X PBS, were individually bubbled with Argon gas for 15 seconds. To prepare 250 µl of click mix we mixed 12.5 µl of 50 mM THPTA, 6.25 µl of 20 mM CuSO_4_, and 2.5 µl of 10 µM dye, and then, the headspace of the 250 µl PCR tube was gently gassed with Argon and incubated for 3 min in the dark. Then, we added 62.5 µl of 100 mM Aminoguanidine, 62.5 µl of 100 mM Sodium Ascorbate, and 104 µl of 1X sterile PBS. The 250 µl of click mix was incubated for 3 more min in the dark and then 42 µl of click mix were added to the PCR tubes containing the 208 µl of sample (previously gassed with Argon). To avoid bubble generation, click mix and sample were not mixed by pipetting. The samples with the click mix were incubated for 30 minutes (Mediterranean samples) or 6 hours (for the Pacific coast and the Mediterranean Sea samples) at room temperature. After the click reaction, samples were concentrated and washed with 0.02-μm-filtered 1X PBS using Amicons (100 KDa, Millipore) to remove unbound dyes as follows. The Mediterranean Sea samples were concentrated one time, washed 3 times with 5 ml 0.02-μm-filtered 1× PBS using 100 KDa 15 ml Amicons (2,500 x g, 3 min), and finally resuspended in 0.02-μm-filtered 1× PBS, 500 µl. The Pacific coast samples were concentrated and washed one time with 500 µl of 0.02-μm-filtered 1X PBS using 0.5 mL Amicons (100 KDa, Millipore) by centrifugation of 7,500 x g, 3 min and resuspended in 0.02-μm-filtered 1× PBS, 500 µl. The Pacific coast sample (500 μl final volume) was additionally counterstained with 2.5 μl 1,000 x SYBR^TM^ Gold, incubated for 20 min in the dark, and washed 3 times with 500 µl of 0.02-μm-filtered 1X PBS in 0.5 mL 100 KDa Amicons (7,500 x g, 3 min). Finally, all samples were immediately analyzed and sorted by FACS within the same day or in less than ≈12 h. In all cases, samples were kept at 4ºC until sorting.

**Fluorescence-activated viral and cell sorting**

FACS analysis and sorting of Pacific coast and Mediterranean samples were conducted in a BD FACSAria^TM^ Fusion (Becton Dickinson, San Jose, CA) at the Caltech Flow Cytometry and Cell Sorting Facility (Pasadena, CA, USA) and Central Service for Experimental Research (SCSIE) from the University of Valencia (Valencia, Spain), respectively. Strict DNA decontamination of instruments, reagents, and materials were performed as described in single cell genomic protocols by Rinke and collaborators^4^ and further studies ^5–8^. In brief, autoclaved sheath fluid was prepared in house with combusted NaCl, sterilized and overnight UV-irradiated. Sheath fluid tank was autoclaved, UV-irradiated, and also decontaminated by washing with 10% bleach, for at least 10 minutes (up to 30 minutes) and rinsed with sterile UV-irradiated (16 h) autoclaved MilliQ water. Then, sheath fluid tank was filled with sterile sheath fluid free of DNA. All this process was carried out on a sterile environment within a PCR hood.

The sorter was setup to detect the red AF647-Pycolyl-Azide (Abs/Em=648/671 nm) fluorescence from BONCAT positive virus and bacteria using the 640 nm red laser. The green fluorescence signal from total (active and non-active) bacteria and DNA-containing viral populations labeled with SYBR^TM^ Gold (Abs/Em=495/537 nm) was excited using the 488 nm blue laser. Both fluorescent channels were represented versus the side scatter height (SSC-H). To delimitate the flow cytometer regions for the different populations, 1X PBS and incubated seawater without HPG were identically processed as samples stained with SYBR^TM^ Gold and BONCAT-click *(Supplementary Fig 2A-C)*. Samples were also additionally filtered by 0.2 μm to better discriminate viral and cell communities (*Supplementary Fig 2D-E)*).

In the cause of Mediterranean samples, the active bacterial and viral progeny populations were sorted based on side scatter height (SSC-H) and APC (BONCAT+) fluorescence intensity (Fig 2). For the Pacific samples, bacterial and viral populations were counterstained with SYBR-Gold as described above and sorted based on the double threshold from FITC (SYBR-Gold) and APC fluorescence (BONCAT) (Figure 2D; Supplementary Figure 2D-E). All sorting experiments were carried out setting the flow cytometry sorter at the single cell mode, which employ the most stringent sorting purity conditions.

Up to five pool replicates of BONCAT-positive populations from Mediterranean samples were sorter in 96-well plates (1,000 and 5,000 viruses from viral progeny; 200 active cells) using a flow rate 40-800 ev/sec. For the Pacific samples, each BONCAT-positive and non-active population was sorted into a single pool in 16h UV irradiated 1.5 ml tubes (100,000 viruses from viral progeny, 50,000 non-active viruses and 25,000 active and non-active bacteria) and divided in wells from 96-well plate with 5,000 viruses or cells each (20 wells for viral progeny, 10 wells for non-active viruses and 5 wells for active and non-active bacteria). Furthermore, 10,000 BONCAT-positive viruses from Pacific Coast were sorter in a 1.5 mL to directly visualize them in the fluorescence microscope (Figure 2D). Overall, during viral sorting, we applied the FACS procedure (e.g. flow rate) previously described in single virus genomic approach^8^ to ensure an efficient sorting of nanoparticles.

The sheath fluid signal detected in the APC channel as electronic noise was also sorted in the same 96-well sterile plates or new sterile 1.5 ml tubes used as a negative control in the multiple displacement amplification (MDA). Sorted cells and viruses were stored at -80 °C until whole-genome amplification.

**Whole-genome amplification of sorted bacterial and viral populations**

The DNA of the viral and cell pools was amplified by MDA using the EquiPhi29 polymerase (ref: A39392; Thermo Fischer scientific)^6^, as described in Garcia-Heredia, et al., 2020^9^. Briefly, after a thermal and chemical lysis of the viral capsid, the MDA master mix was added containing 0.26 U of Equiphi29 DNA polymerase (Thermo Fischer scientific), 1X EquiPhi29 reaction buffer (Thermo Fischer scientific), 0.8 µl of 0.04 mM heptamers (IDT), 10 mM of DTT (Sigma), 0.4 mM of dNTPs (New England Biolab), 0.002 µl of SYTO 9 (Invitrogen) and 5.44 µl of sterile ultraviolet 16 h-treated milliQ water. During the preparation, MDA master mix, except SYTO 9, and the DLB (0.4 M KOH, 10 mM EDTA and 100 mM dithiothreitol) and stop solution (pH=4, Qiagen) used for viral/cell lysis were UV-light irradiated as described in Rinke et al., 2014^4^. The final MDA reaction volume was scaled upon the volume of sorted samples. As MDA positive controls, we added 1 ng/µl and 10 ng/ µl genomic DNA for the *E.coli* lambda phage, and 16 h-UV-irradiated 1X TE buffer as negative control. The real-time monitored reaction of MDA was carried out at 45ºC. All sample wells from the Pacific Coast samples amplified, while 77 % of the sample wells from the Mediterranean Sea resulted in successful amplification. None of the wells containing sorted sheath fluid amplified confirming the correct DNA decontamination procedure and cleaning process. The MDA reactions were stopped at 75ºC for 10 minutes and the plates were stored at -80ºC until use.

**Sequencing and assembly of amplified genome from viral and bacterial populations**

All the MDA products from the Pacific Coast samples were sequenced (100,000 BONCAT-positive viruses, 50,000 non-active viruses, and 25,000 active and non-active bacteria). A subset of 5 pools were selected from the Mediterranean Sea for sequencing; 3 from the 30 min click reaction (1 x 200 active cells and 1 x 1,000 and 1 x 5,000 BONCAT-positive viruses) and 2 from 6h click reaction (1 x 200 and 1 x 5,000 active cells and BONCAT-positive viruses respectively). Libraries were prepared using the Illumina DNA Prep (M) kit and adapters from IDT for Illumina DNA/RNA UD, following the manufacturer’s protocol. Illumina paired-end sequencing was performed with NovaSeq 6000 (150 x 2 PE; approximately 3 Gb per sample) by Macrogen (Korea). The obtained sequence reads were quality-trimmed using Trimmomatic 0.39^10^ with the following settings: -phred33 LEADING:3 TRAILING:3 SLIDINGWINDOW:4:30 MINLEN:50. Quality-filtered reads were assembled using Spades v3.15.5^11^ with the following settings: --sc mode, -k 21,33,55,77,99,127.

**Taxonomic analysis of bacterial reads**

Trimmed and unassembled reads derived from both active and non-active bacterial samples were taxonomic analyzed using Kaiju^12^ , and compared against the NCBI BLAST database (nr_euk) which includes data from bacteria, archaea, eukaryotes and viruses, with the following parameters: -E 0.00001, -a greedy. A taxonomic profiling with krona^13^ was generated and analyzed.

**Viral identification and bioinformatic genome analysis**

For detecting and classifying viral contigs, after assembly, a combination of Virsorter2.0 (v2.2.3)^14^ and CheckV (v0.9.0)^15^ (following the viral sequence identification SOP V.3 from the Sullivan Lab, **[dx.doi.org/10.17504/protocols.io.bwm5pc86](https://dx.doi.org/10.17504/protocols.io.bwm5pc86" \t "_blank)**), and geNomad (v1.7.0; default parameters)^16^ was used. In addition, to corroborate the viral origin of the detected viral contigs, and better characterize some specific viral groups, the ORFs from all contigs ≥ 1,500 bp were predicted using Prodigal v2.6.3^17^ and compared against the high confidence viral proteins from the IMG/VR v4^18^ with Protein-Protein BLAST 2.8.1+ using the following settings: *-outfmt "6 qseqid sseqid pident length mismatch gapopen qstart qend sstart send evalue bitscore qlen slen positive ppos", -evalue 0.00001*. We considered as *bona fide* viruses all these genomes detected by the viral identification SOP, also by geNomad and sharing more than 2 proteins (thresholds of 80% similarity and a coverage of 90%) with viral contigs from IMG/VR.
To search OM43 phages within our viral contigs we employed as reference the isolated OM43 phages availables from NCBI, the phage Venkman EXVC282S^19^, the Methylophilales phage MEP301^20^, the Methylophilales phages Melnitz (EXVC039-EXVC044M)^21^ , the Methylophilales phages MEP401-MEP402 and their related 99 metagenomic viral genomes found in viral databases ^22^.

The reference genome of the vSAG 37-F6^8,23^, considered one of the most abundant and widespread marine viruses, was also used to search similar viruses within that population. Following the same protocol described above, their proteins were blastp compared with all the viral-BONCAT genomes, finding 2 viral contigs sharing over 16 viral proteins, with an average amino acid similarity of approximately 74-75%.

To polish the taxonomic annotation, those contigs > 5,000 bp detected with geNomad from all samples as “giant” viruses and virophages (Phylum Nucleocytoviricota and Preplasmiviricota, respectively) were manually verified with the taxonomic information from the IMG/VR metadata.

**Viral protein network**

To build a network clustering similar viruses based on protein sharing we employed vConTACT2 (v0.9.19)^24^ through the DOE Systems Biology Knowledgebase (KBase, [http://kbase.us](http://kbase.us/))^25^ using as reference database the “Prokaryotic Viral RefSeq 201”, protein cluster method “MCL” and viral cluster method “ClusterONE”. The viral network was visualized using Cytoscape v3.7.1^26^

As an indicator of the viral turnover rate of each viral population we estimated the relative viral progeny ratio as the percentage of viral BONCAT positive viruses over the total viruses (active and inactive) from a cluster (e.g. Fig. 4 and Supplementary Table 1). As only inactive viruses from the Pacific Ocean were sorted and analyzed; calculations of viral progeny ratio only used datasets from the Pacific Ocean viral genomes associated with the BONCAT positive and negative-sorted free virions.To account for differences in sequencing depth caused by the initial sorting strategy (which included 100,000 viral progeny viruses but only 50,000 inactive viruses) we normalized the data using the formula:

(1) Viral progeny ratio (%) = 100 x number of active viruses (viral progeny)/ (number of active + 2 x number of inactive viruses).

This correction reflects that approximately twice as many viral genomes were retrieved from viral progeny samples (n=543) as from inactive ones (287), thereby avoiding overestimation of active viruses in the dataset (detailed calculations of the viral progeny ratio for each cluster, along with counts of active and inactive viruses, are provided in Supplementary Data 1).

**Far-T4 phages identification**

For the Far-T4 phages identification we used an approximation like that used by Roux, S., *et al*., 2015 ^27^, screening all the viral BONCAT contigs > 1,500 bp for the presence of the T4 major capsid protein Gp23. All Gp23-like protein (protein to protein blastp bit score > 50 and e-value < 1x10^-10^) were selected to build a phylogenetic tree. For it, all the Gp23-like proteins were aligned using MAFFT v7.310 (--auto)^28^, columns with gaps in more than 80% of the sequences were removed from the alignment using trimAI v1.4 (-gt 0.2)^29^ and then the tree was constructed using Geneious bioinformatic software^30^. In parallel, all the proteins from our viral contigs were blastp-compared against the proteins of *Rhodothermus RM378 phage* (as the only isolated representative of the Far-T4 phages group) to check the presence of core proteins.

**Host assignment of the Far-T4 phage cluster**

To computationally identify the hosts of the viruses contained the Far-T4 phage cluster, we run the iPHoP machine learning (v1.3.3)^31^ using the host database “*iPHoP_db_Aug23_rw*” based on GTDB r214 and extended collection of MAGs. Due to that application did not show significant results (probably due to the completeness percentage of our sequences), we opted for an indirect approach; we compared all the proteins of the viruses from the Far-T4 phages cluster with those of the high confident IMG/VR v4 database^18^ using blastp. To be conservative, we selected those IMG/VR genomes sharing at least the 80% of the proteins of our contigs (not less than 5 proteins) with > 60% of amino acid similarity and >95% amino acid coverage of the alignment. Then iPHoP was applied to that IMG/VR contigs as described above.

**Bacterial read fragment recruitment against OM43 genomes**

To estimate the abundance of the OM43 clade in active and non-active bacterial fraction from Pacific and Mediterranean seawater, we performed a read recruitment plot analysis^32^ using 13 publicly available genomes at NCBI derived from isolates OM43 HTCC2181^33^, KB13^34^, MBRSH7^35^, H5P1^21^and LSUCC0622, LSUCC0268, LSUCC0389, LSUCC0401, LSUCC0536, LSUCC0568, LSUCC0603, LSUCC0665, LSUCC0717^36^. Trimmed reads derived from bacterial samples were subjected to BLASTn analysis (BLAST 2.8.1+) against concatenated OM43 genomes using the following settings: blastn -outfmt "6 qseqid sseqid pident length mismatch gapopen qstart qend sstart send evalue bitscore qlen slen ", -evalue 0.00001, and applying a threshold of 70% similarity and a coverage of 70%. The BLASTn output was filtered using the BestHit Script available by enveomics tools^32^ and recruitment plots were carried out using enve.recplot 2 (R stadistic pluging).

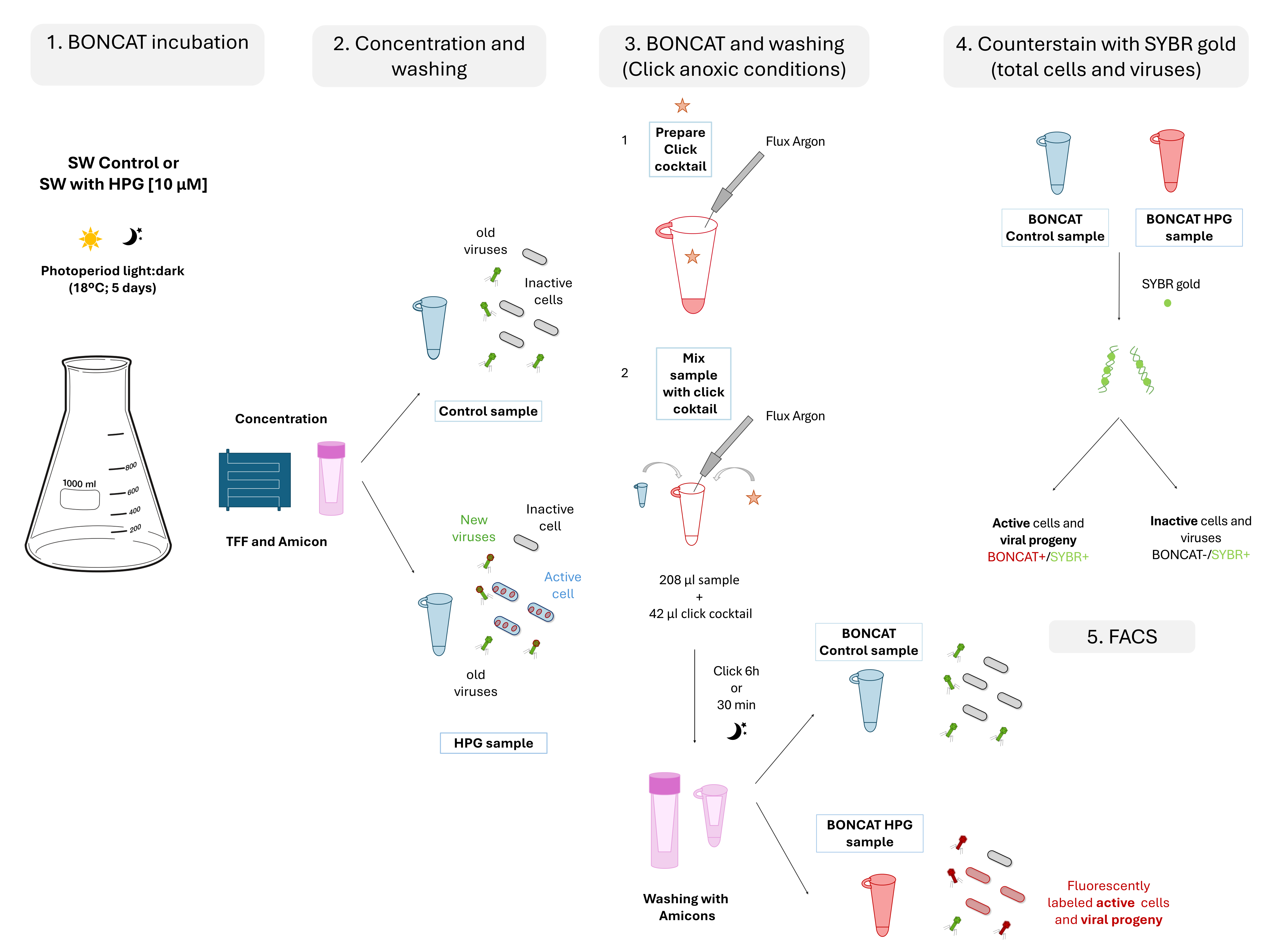

**Supplementary Figure 1**. Scheme of the BONCAT-FACS workflow for seawater. Briefly, control and HPG samples (amended with 10 µM HPG) were incubated for 5 days with light:dark photoperiods. Sample concentration and washing was carried out by tangential flow filtration (TFF) and 100 KDa Amicons, respectively. After preparing the click mix in 2 steps (1), the BONCAT reaction will begin by mixing 208 µ of sample with 42 μl of click mix in 250 μl tubes (2), minimizing the available air space and insufflating with argon. After the BONCAT reaction (6 h or 30 min), washing steps were included to remove unbound dyes. BONCAT Mediterranean coast samples were analyzed and sorted directly with FACS (BONCAT + cells and viruses), while Pacific coast samples were counterstained with SYBR Gold and sorted with double threshold (BONCAT +/SYBR + cells and viruses; BONCAT -/SYBR + inactive cells and viruses).

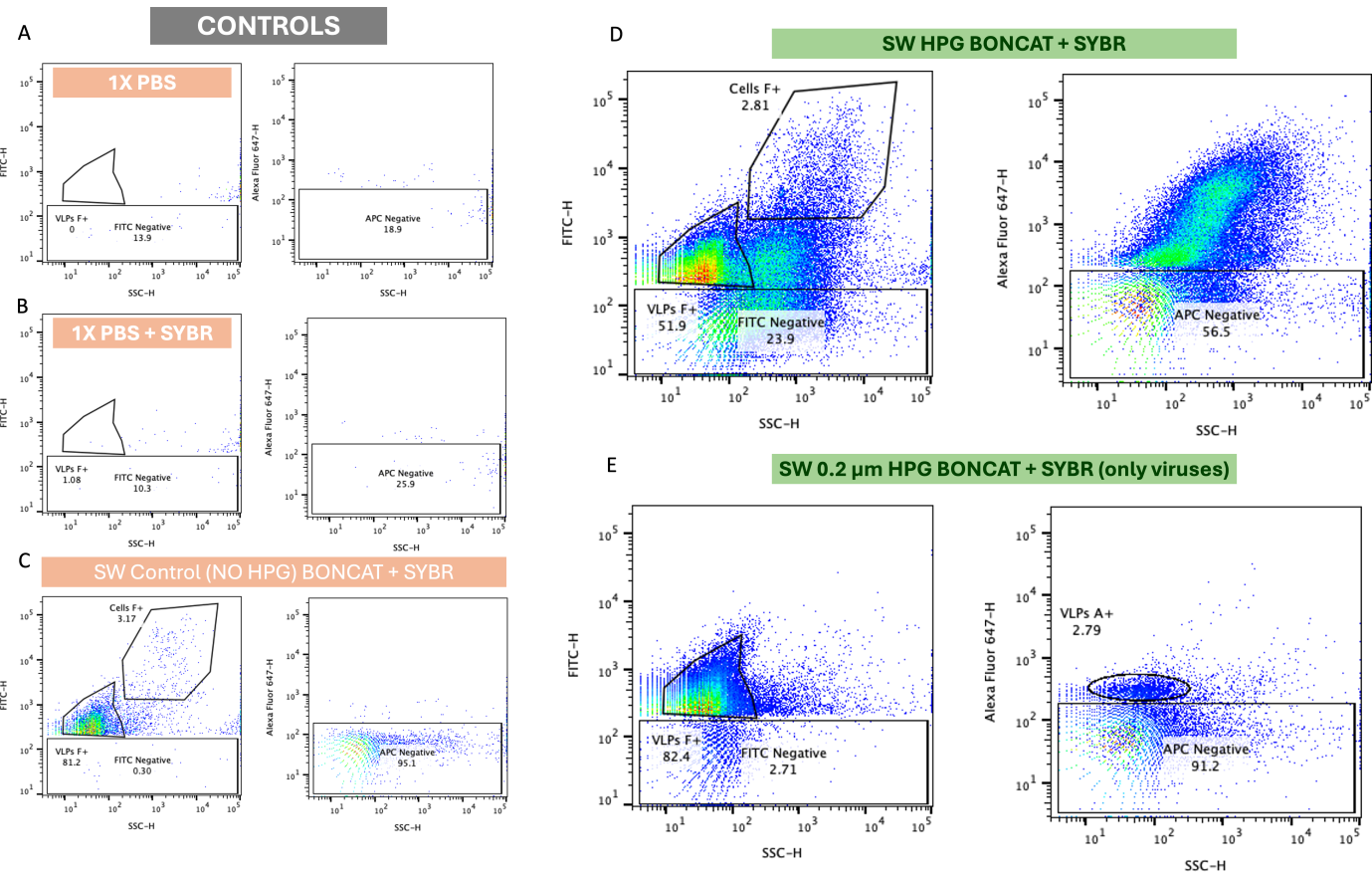

**Supplementary Figure 2**. Flow cytometry plots showing gates used for sorting the viral progeny and active cell populations (BONCAT +). Two different fluorescent channels were employed, the green one to visualize the SYBR-Gold DNA staining (total cells and viruses) and the red one to detect the AF647-Pycolyl-Azide dye used for staining and discriminating the BONCAT + active cells and viruses (HPG incubated). Controls A-C were used as negative samples containing PBS buffer (used to wash and resuspend the samples) with and without SYBR-Gold, and the sample incubated without HPG after the click reaction. These controls are necessary to discriminate the noise background vs sample signal. D) Flow cytometry plots corresponding to the doubled stained sample. The viral and cell populations delimited in the SYBR-Gold channel was observed. E) Finally, by filtering the sample by 0.2 μm, active cells were removed, and the active viral progeny fraction was the only one observed, which allowed us to discriminate the gate for sorting of viral progeny.

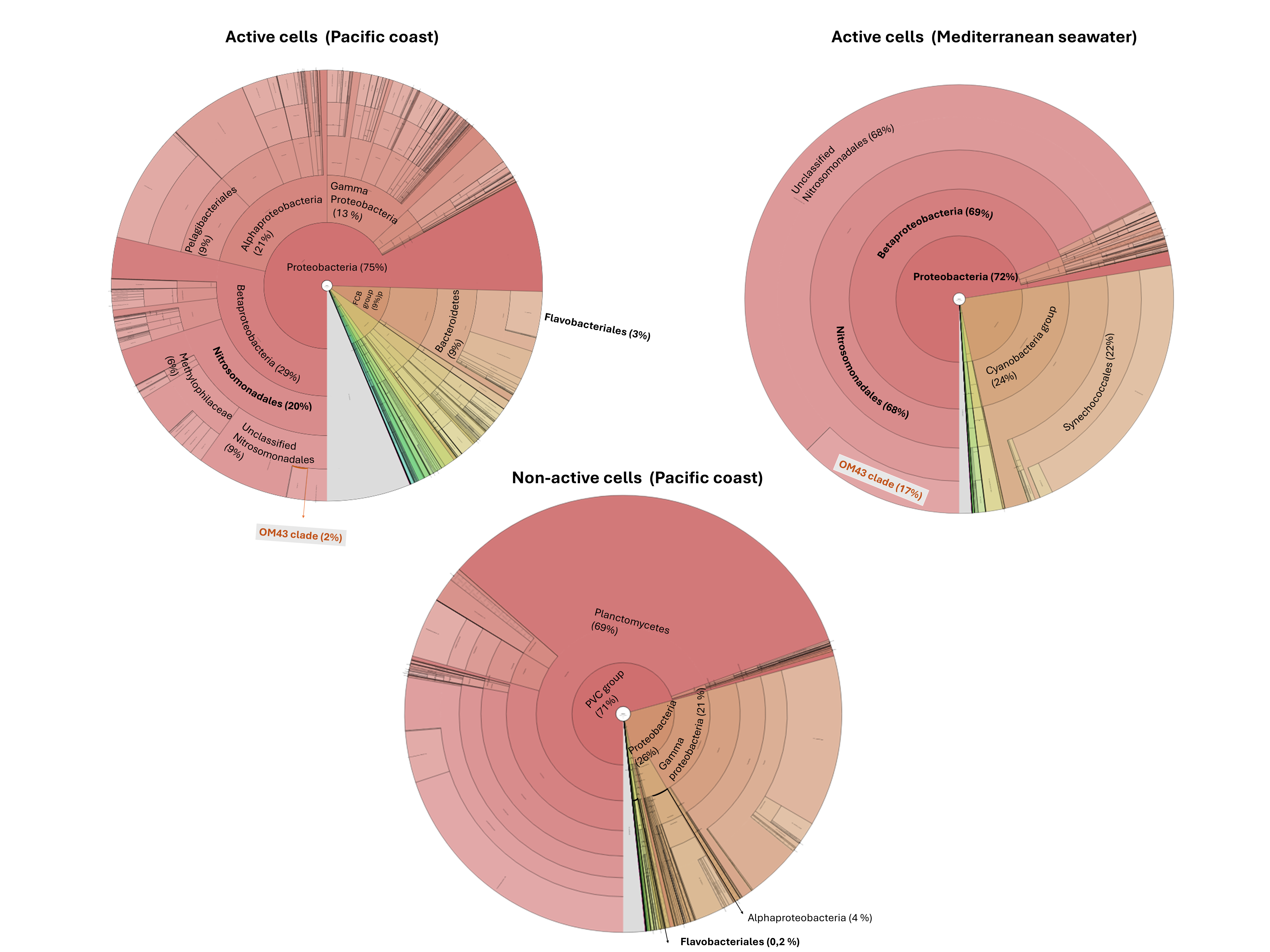

**Supplementary Figure 3**. Taxonomic assignment of sequenced reads obtained from the active and non-active cell fractions. Analysis was performed with Kaiju program (see methods for details). The most abundant bacterial groups and their relative abundances (expressed as percent of total bacterial reads) are highlighted in bold in the taxonomic graphs. The relative abundance of the OM43 clade is shown as a percentage of total proteobacteria reads.

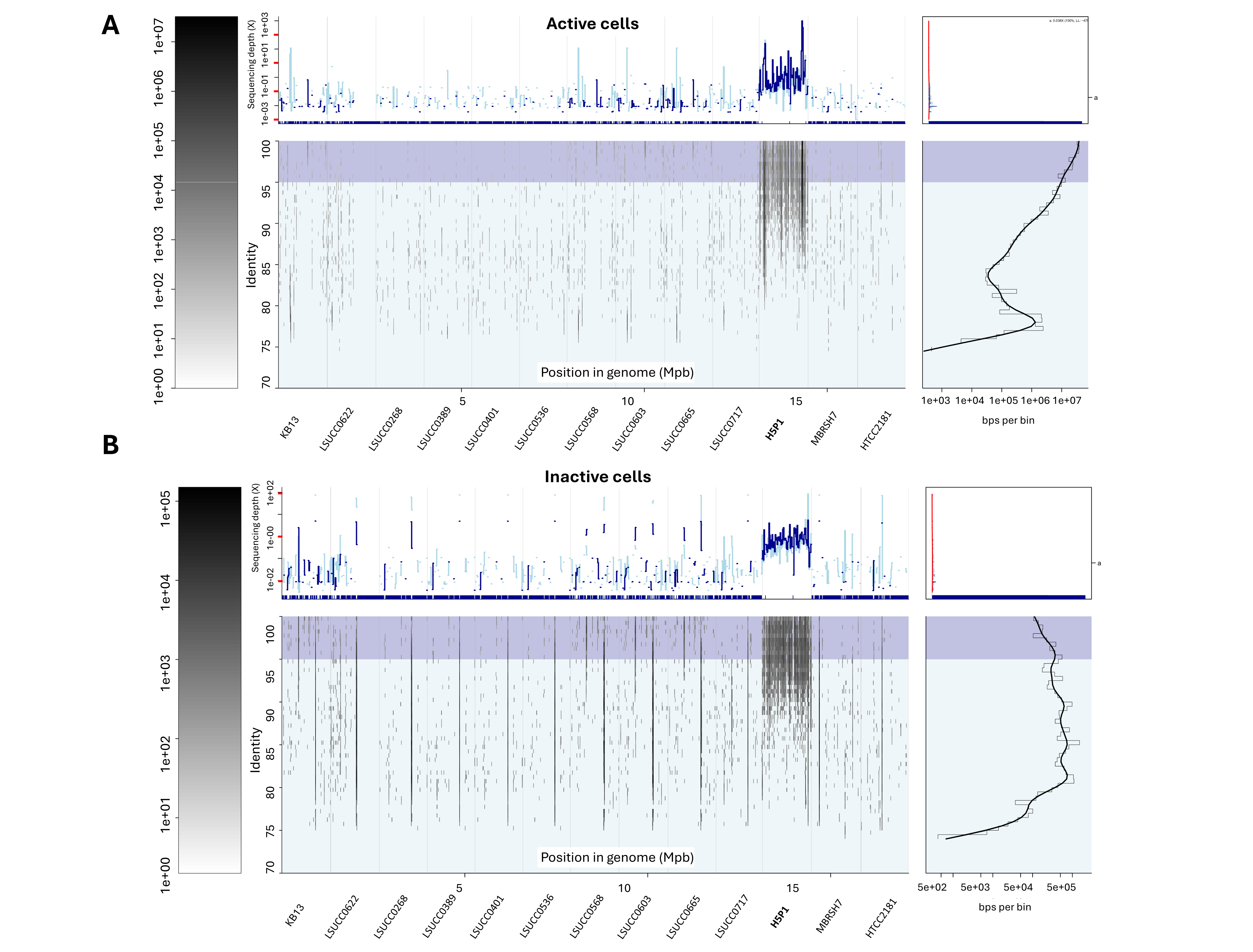

**Supplementary Figure 4**. Fragment recruitment of the active (A) and non-active cell (B) reads from the Pacific Ocean against 13 different OM43 genomes from isolated strains. Strain *Methylophilales bacterium* H5P1 (marked in bold) was recovered in both types of samples.

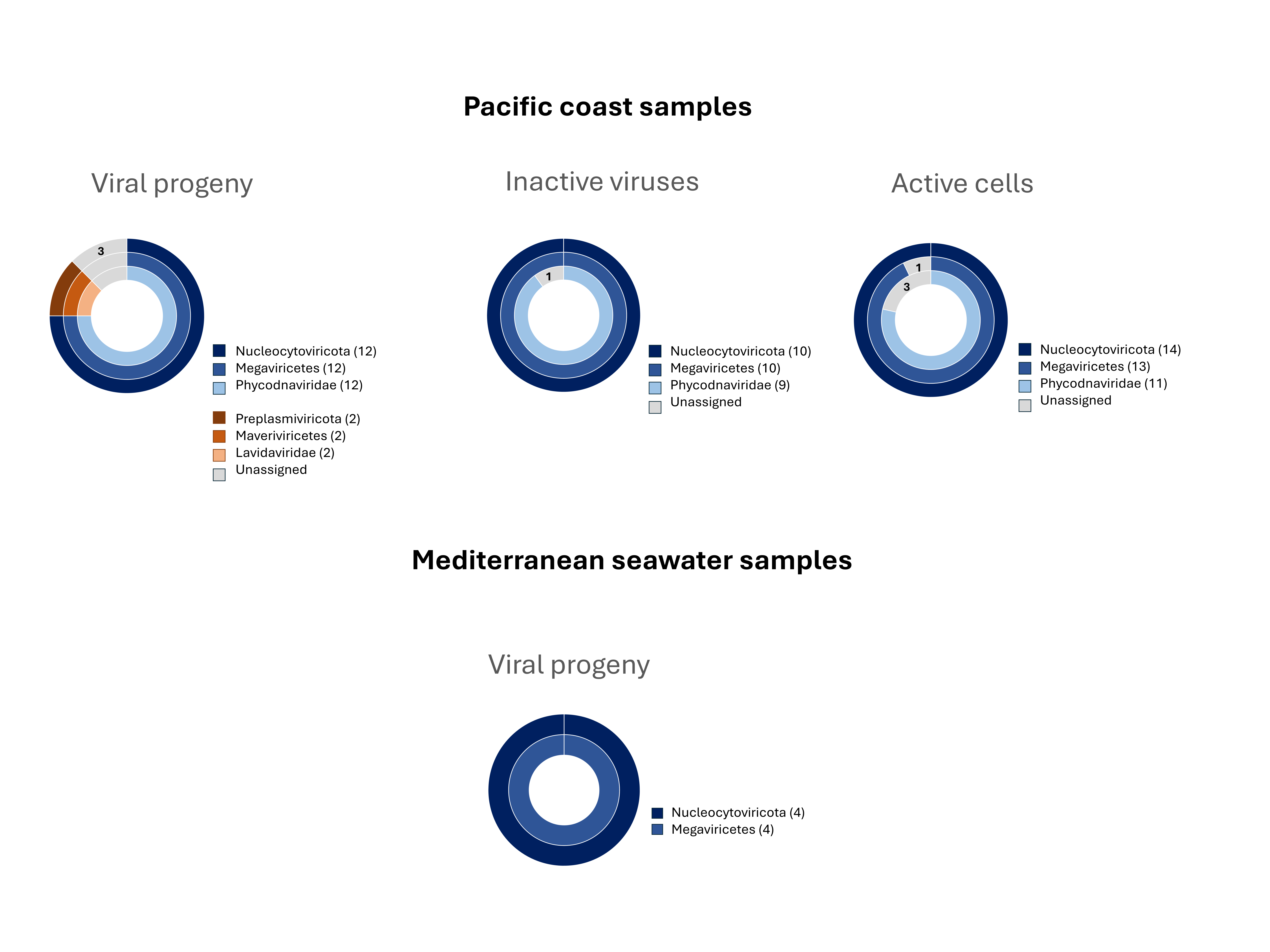

**Supplementary Figure 5.** Eukaryote viral contigs recovered in the Pacific and the Mediterranean coastal samples identified as large double stranded DNA viruses within the 40 Nucleocytoviricota and putative Nucleocitoviricota-infecting Virophages from the different BONCAT fractions. A combination of geNomad analysis and BLASTp against IMG/VR 4.1 database was used to corroborate these taxonomic assignments. Contigs without a definitive taxonomic assignment are listed as unassigned.

**Supplementary Table 1:** Viral progeny ratio of the clusters identified in the viral network shown in Figure 4. The table shows the number of viruses identified as viral progeny (VP) and inactive viruses (VI) within each cluster. The viral progeny ratio was calculated using the formula (1) to account for differences in sequencing depth due to the initial sorting strategy. Only datasets from Pacific Ocean samples were included, as both inactive and active fractions were sorted.

| **Cluster** | **Active viruses (VP)** | **Inactive viruses (VI)** | **n**  **(active + inactive)** | **viral progeny ratio** |
| --- | --- | --- | --- | --- |
| Cluster Far-T4-like phages | 180 | 19 | 199 | 82.57 |
| Cluster 37-F6-like phages | 15 | 2 | 17 | 78.95 |
| Cluster 1 | 68 | 66 | 134 | 34.00 |
| Cluster 2 | 17 | 27 | 44 | 23.94 |
| Cluster 3 | 43 | 1 | 44 | 95.56 |
| Cluster 4 | 62 | 16 | 78 | 65.96 |
| Cluster 5 | 93 | 41 | 134 | 53.14 |

**Supplementary Table 2.** Active (viral progeny) and non-active viral contigs associated with the large double stranded DNA viruses, Nucleocytoviricota according to geNomad assignment. These putative eukaryote-infecting viruses were also recovered from the active cell fraction. Data was later corroborated by viral protein comparison against the high-confidence IMG/VR v4.1 viral database. Contigs detected as Mimiviridae by geNomad but without classification assignment by BLASTP are shown in blue.

| **Sample** | **Viral** | **Population^1^** | **Length** | **Virus** | **Taxonomy (geNomad assignation)** | **IMG/VR** | **IMG/VR** | **IMG/VR** | **IMG/VR** |
| --- | --- | --- | --- | --- | --- | --- | --- | --- | --- |
|  | **contig** |  |  | **Score^2^** |  | **Hits^3^** | **Hits taxonomy^4^**  **(Nucleoviricota)** | **Hits Taxonomy^4^ (Megaviricetes)** | **Hits Taxonomy^4^ (Family)** |
| Pacific Ocean | Contig 164 | VP | 8,390 | 0,9831 | Megaviricetes;Algavirales;Phycodnaviridae | 362 | 362 | 358 | 356 |
|  | Contig 182 | VP | 8,121 | 0,9736 | Megaviricetes;Algavirales;Phycodnaviridae | 283 | 283 | 282 | 280 |
|  | Contig 232 | VP | 7,363 | 0,9764 | Megaviricetes;Algavirales;Phycodnaviridae | 97 | 97 | 97 | 96 |
|  | Contig 234 | VP | 7,346 | 0,9828 | Megaviricetes;Algavirales;Phycodnaviridae | 176 | 176 | 176 | 175 |
|  | Contig 275* | VP | 6,978 | 0,9738 | Megaviricetes;Algavirales;Phycodnaviridae | 3 |  |  |  |
|  | Contig 809 | VP | 6,921 | 0,9672 | Megaviricetes;Algavirales;Phycodnaviridae | 343 | 343 | 343 | 341 |
|  | Contig 839 | VP | 6,781 | 0,9761 | Megaviricetes;Algavirales;Phycodnaviridae | 588 | 588 | 565 | 565 |
|  | Contig 316 | VP | 6,419 | 0,9799 | Megaviricetes;Algavirales;Phycodnaviridae | 53 | 53 | 53 | 53 |
|  | Contig 321 | VP | 6,361 | 0,9822 | Megaviricetes;Algavirales;Phycodnaviridae | 154 | 154 | 154 | 152 |
|  | Contig 986 | VP | 5,936 | 0,9824 | Megaviricetes;Algavirales;Phycodnaviridae | 420 | 420 | 419 | 417 |
|  | Contig 993 | VP | 5,893 | 0,9671 | Megaviricetes;Algavirales;Phycodnaviridae | 126 | 123 | 123 | 121 |
|  | Contig 1057 | VP | 5,512 | 0,9793 | Megaviricetes;Algavirales;Phycodnaviridae | 97 | 97 | 97 | 93 |
|  | Contig 396 | VP | 5,386 | 0,9821 | Megaviricetes;Algavirales;Phycodnaviridae | 201 | 201 | 201 | 198 |
|  | Contig 148 | VI | 11,660 | 0,9796 | Megaviricetes;Algavirales;Phycodnaviridae | 325 | 325 | 325 | 316 |
|  | Contig 188 | VI | 10,456 | 0,9812 | Megaviricetes;Algavirales;Phycodnaviridae | 183 | 182 | 182 | 173 |
|  | Contig 194 | VI | 10,355 | 0,9818 | Megaviricetes;Algavirales;Phycodnaviridae | 90 | 90 | 90 | 90 |
|  | Contig 332 | VI | 7,413 | 0,9697 | Megaviricetes;Algavirales;Phycodnaviridae | 63 | 63 | 63 | 61 |
|  | Contig 340 | VI | 7,237 | 0,9817 | Megaviricetes;Algavirales;Phycodnaviridae | 370 | 370 | 370 | 359 |
|  | Contig 425 | VI | 5,967 | 0,9180 | Megaviricetes;Algavirales;Phycodnaviridae | 10 | 10 | 10 | 10 |
|  | Contig 453 | VI | 5,546 | 0,9781 | Megaviricetes;Algavirales;Phycodnaviridae | 21 | 21 | 21 | 15 |
|  | Contig 479 | VI | 5,276 | 0,9815 | Megaviricetes;Algavirales;Phycodnaviridae | 79 | 79 | 79 | 78 |
|  | Contig 491 | VI | 5,110 | 0,9607 | Megaviricetes;Algavirales;Phycodnaviridae | 83 | 83 | 83 | 82 |
|  | Contig 175 | VI | 10,882 | 0,9815 | Megaviricetes;Imitervirales;Mimiviridae | 4 | 4 | 4 |  |
|  | Contig 38* | CA | 12,336 | 0,9785 | Nucleocytoviricota;unassigned | 5 |  |  |  |
|  | Contig 292 | CA | 6,117 | 0,9757 | Megaviricetes | 4 | 3 | 3 | 3 |
|  | Contig 284 | CA | 6,237 | 0,9830 | Megaviricetes;Algavirales;Phycodnaviridae | 281 | 279 | 279 | 278 |
|  | Contig 264 | CA | 6,468 | 0,9824 | Megaviricetes;Algavirales;Phycodnaviridae | 150 | 150 | 150 | 149 |
|  | Contig 265 | CA | 6,467 | 0,9819 | Megaviricetes;Algavirales;Phycodnaviridae | 17 | 17 | 17 | 16 |
|  | Contig 113 | CA | 8,999 | 0,9818 | Megaviricetes;Algavirales;Phycodnaviridae | 557 | 557 | 557 | 550 |
|  | Contig 167 | CA | 7,868 | 0,9816 | Megaviricetes;Algavirales;Phycodnaviridae | 227 | 227 | 223 | 219 |
|  | Contig 169 | CA | 7,826 | 0,9806 | Megaviricetes;Algavirales;Phycodnaviridae | 583 | 583 | 580 | 574 |
|  | Contig 210 | CA | 7,116 | 0,9799 | Megaviricetes;Algavirales;Phycodnaviridae | 451 | 451 | 451 | 447 |
|  | Contig 336 | CA | 5,629 | 0,9797 | Megaviricetes;Algavirales;Phycodnaviridae | 358 | 358 | 357 | 357 |
|  | Contig 350 | CA | 5,501 | 0,9736 | Megaviricetes;Algavirales;Phycodnaviridae | 447 | 447 | 447 | 422 |
|  | Contig 32 | CA | 13,324 | 0,9712 | Megaviricetes;Algavirales;Phycodnaviridae | 367 | 367 | 367 | 359 |
|  | Contig 186 | CA | 7,495 | 0,9497 | Megaviricetes;Algavirales;Phycodnaviridae | 236 | 236 | 236 | 233 |
|  | Contig 78 | CA | 10,051 | 0,7260 | Megaviricetes;Imitervirales;Mimiviridae | 1 | 1 | 1 | 1 |
| Mediterranean Sea | Contig 12 | VP | 16,033 | 0,9735 | Megaviricetes;Algavirales;Phycodnaviridae | 2 | 2 | 2 |  |
|  | Contig 13 | VP | 15,711 | 0,9726 | Megaviricetes;Pimascovirales;Iridoviridae | 4 | 4 | 4 |  |
|  | Contig 7 | VP | 26,621 | 0,9724 | Megaviricetes | 3 | 3 | 3 |  |
|  | Contig 3 | VP | 67,645 | 0,9071 | Megaviricetes | 2 | 2 | 2 |  |

1 Population. Viral genomes obtained from the different sorted populations, CA: Active cells, VP: Viral progeny, VI : Non-active viruses.

2 Virus-score. Obtained from geNomad viral identification

3 IMG/VR hits: Number of hits (80% Similarity, 90% hit coverage) with high-confident viral genomes from IMG/VR v4.1

4 IMG/VR Hits taxonomy: Number of hits with the same taxonomic level assignment compared to geNomad

* contigs hits with Bamfordvirae kingdom according to IMG/VR v4.1

**Supplementary Table 3.** Detection of small double stranded DNA virophages Active (viral progeny, VP) and non-active viral contigs (VI) from Preplasmiviricota according to geNomad assignment. Data was later corroborated by viral protein comparison against the high-confidence IMG/VR v4.1 viral database.

| **Sample** | **Viral** | **Population^1^** | **Length** | **Virus** | **Taxonomy (geNomad)** | **IMG/VR** | **IMG/VR** | **IMG/VR** | **IMG/VR** |
| --- | --- | --- | --- | --- | --- | --- | --- | --- | --- |
|  | **contig** |  |  | **Score^2^** |  | **Hits^3^** | **Hits taxonomy^4^**  **(Preplasmoviricota)** | **Hits Taxonomy^4^ (Maveriviricetes)** | **Hits Taxonomy^4^ (Family)** |
| Pacific Ocean | Contig 29* | VP | 14,029 | 0,9755 | Maveriviricetes;Priklausovirales;Lavidaviridae | 5 |  |  |  |
|  | Contig 217 | VP | 7,606 | 0,9816 | Maveriviricetes;Priklausovirales;Lavidaviridae | 9 | 6 | 5 | 5 |
|  | Contig 1012 | VP | 5,760 | 0,9759 | Maveriviricetes;Priklausovirales;Lavidaviridae | 177 | 94 | 81 | 27 |
|  | Contig 239* | VI | 9,134 | 0,9722 | Maveriviricetes;Priklausovirales;Lavidaviridae | 2 |  |  |  |

1 Population. Viral genomes obtained from the different sorted populations, VP: Viral progeny, VI : Non-active viruses.

2 Virus-score. Obtained from geNomad viral identification

3 IMG/VR hits: Number of hits (80% Similarity, 90% hit coverage) with high-confident viral genomes from IMG/VR v4.1

4 IMG/VR Hits taxonomy: Number with the same level of taxonomic assignment compared to geNomad

* contigs hits with Bamfordvirae kingdom according to IMG/VR v4.1

**Supplementary Data 1-8 legends**

**Supplementary Data 1. Viral progeny ratio per cluster in the viral network**

Detailed calculation of the viral progeny ratio for the clusters identified in the viral network. This table includes all viral contigs assigned to each of the seven clusters identified in the network. For each cluster, contigs are labeled as *reference* if they correspond to reference genomes from the vContact2 database, *VP* if they belong to the active (viral progeny) fraction, and *VI* if they belong to the inactive viral fraction. The value of *n* reflects the number of viral contigs from our samples (active + inactive) detected within each cluster. Clusters highlighted in red exhibit a high viral progeny ratio, while those in blue show a low viral progeny ratio.

**Supplementary Data 2. Similarity of viral-BONCAT contigs to vSAG 37-F6 proteins**

Results of a BLASTp search using ORFs predicted from the vSAG 37-F6 genome (Prodigal v2.6.3) against all viral-BONCAT contigs. Two contigs share >16 proteins with vSAG 37-F6, with ~74–75% average amino acid identity.

**Supplementary Data 3. Predicted microbial hosts of Far-T4 phages**

Host predictions based on iPHoP using IMG/VR v4 genomes sharing ≥80% of proteins with Far-T4 phages. Flavobacteriales-affiliated genomes are highlighted in green**.**

**Supplementary Data 4. Taxonomic profile of bacterial samples at the order level**

Taxonomic classification using Kaiju against the NCBI nr_euk database. The table is collapsed at the order level, showing relative abundances as percentage of total reads (percent_total) and percentage of bacterial reads only (percent_bacteria). Flavobacteriales are highlighted in green; the most active bacterial orders in blue.

**Supplementary Data 5. Taxonomic profile at the phylum level**

Taxonomic classification using Kaiju against the NCBI nr_euk database. The table is collapsed at the phylum level. The most abundant phylum in active fractions is highlighted in blue; in inactive fractions, in red.

**Supplementary Data 6. Taxonomic profile at the family level**

Taxonomic classification using Kaiju against the NCBI nr_euk database. The table is collapsed at the family level. The most abundant bacterial family in the inactive fractions is highlighted in red.

**Supplementary Data 7. Taxonomic profile at the species level**

Taxonomic classification using Kaiju against the NCBI nr_euk database. The table is collapsed at the species level. For species in the phylum Proteobacteria (formerly Pseudomonodota), an additional column shows their percentage within the phylum (percent_Pseudomonadota). OM43 clade species are highlighted in blue.

**Supplementary Data 8.OM43 phage identification**

BLASTp comparison of viral-BONCAT contigs against known OM43 phages (e.g., Melnitz, Venkman, MEP301, MEP401–402) and 99 related metagenomic viral genomes. Four contigs were closely related to Melnitz, known to infect OM43 strain H5P1.
